# Good parenting of oil-collecting bees: microbial defense in nests of Centris bees?

**DOI:** 10.64898/2026.03.23.713567

**Authors:** Elif Kardas, Miguel Pacheco-Leiva, Allan Artavia-León, Mauricio Fernández Otarola, Gabriel Vargas Asensio, James D. Ackerman, Adrián A. Pinto-Tomás, Filipa Godoy-Vitorino

## Abstract

Microbial symbionts are increasingly recognized as key contributors to bee health, yet their roles in solitary bee brood cells remain largely unexplored. Here, we characterize the bacterial and fungal communities associated with brood cell compartments (cocoon, meconium, and prepupa) of the oil-collecting bee *Centris aethyctera*, and compare them with gut microbiota from adult *Centris* species. Using 16S rRNA and ITS amplicon sequencing, we show that microbiota are strongly compartmentalized, with distinct diversity patterns, taxonomic compositions, and inferred functional profiles across brood cell components. Contrary to our initial hypothesis, antibiotic-producing bacteria, particularly Actinomycetia, are most diverse and abundant in the meconium rather than the cocoon. Cocoons are enriched in hydrocarbon-degrading and nitrogen-cycling bacteria, while pre-pupae harbor distinct bacterial and fungal taxa, including genera with potential antimicrobial and symbiotic functions. Fungal communities are likewise structured, with taxa such as *Aspergillus* and *Lecanicillium* suggesting possible roles in pathogen defense. Core gut microbiota of adult *Centris* include acetic acid bacteria shared across species, with partial overlap with brood cell taxa, indicating potential transmission pathways. Together, our results reveal that *Centris* brood cells form a highly structured, antimicrobial-rich microenvironment likely shaped by maternal provisioning and environmental acquisition. These findings provide the first comprehensive description of microbiota across brood cell compartments in a solitary bee and identify ground-nesting bee systems as promising reservoirs for novel antimicrobial discovery.

## Introduction

Evidence is accumulating on the importance of bacteria and fungi in pollen and nectar provisions of bees. These are essential for larval development [1–3], immune response, resistance to disease [4,5], and overall bee fitness [6–9]. Microbes present in the bee nest originate from the environment, e.g., plant and nest material, pollen or nectar [6,10–14]. The microbiota can either be transmitted vertically or horizontally. The former refers to the microbial transmission from the mother bee to the egg through the reproductive tract. The latter refers to the microbial transmission between individuals of the same colony, between individuals from different colonies, from the mother bee digestive tract, through specialized structures to carry them, or directly retrieved from the environment, e.g., pollen, nectar or oil [12,15].

Insects in general use symbiotic antibiotic-producing bacteria as a defense line during reproduction [16–22]. For instance, some female herbivorous beetles (Tenebrionidae: Lagriinae) have antibiotic-producing symbionts, *Burkolderia gladioli*, in glands connected to their reproductive system. They deposit these onto the surface of their eggs as a defense against an entomopathogenic fungus, *Purpureocillium lilacinum* [16]. Antibiotic-producing symbionts occur in Hymenoptera as well, especially for immatures and their food provisions in groundnesting species, as they are vulnerable to attack by soil-borne microbes. Fungi-growing ants (Formicidae: Attini) use bacteria, *Pseudonocardia* spp., which produce antibiotics to protect themselves and their fungal gardens against parasitic fungi [19,23–25]. Solitary wasps also have actinomycetes in their nests, such as *Streptomyces*, *Micromonospora*, and *Actinoplanes* which produce important bioactive secondary metabolites like antibiotics[26]. In particular, “*Candidatus Streptomyces philanthi”* protects cells of beewolf wasps (Crabronidae: *Philanthus*) against fungal infestations. The wasp cultivates the actinomycete in specialized antennal glands and applies them to the brood cell prior to oviposition [20,27–29]. In fact, some immature Hymenoptera use an antibiotic mixture against fungal and bacterial antagonists which slows down evolution of resistance [30].

In contrast, antimicrobial use in bees is seldom reported, [e.g., 31,32]. In honeybee hives the greatest abundance of actinomycetes occurs in stored pollen. The same antimicrobialproducing bacteria are also present in older adult bees (hive bees, foraging bees, swarming bees or commercial bees), rather than in the bee brood or hive environment (pupae, newly eclosed workers, honey, empty combs, or propolis) [33].

The nests of solitary bees also have actinomycetes. *Nocardioides* spp. and *Streptomyces* spp. occur in the oil-collecting *Centris decolorata* (Apidae) from Puerto Rico, specifically in the mother bee gut. Sequencing of 16S rRNA markers from the gut of *Centris decolorata*, highlighted the presence of an uncultured *Streptomyces* in various samples potentially representing a new family of *Streptomyces*-derived antibiotics [34].

To date, the diversity and composition of antibiotic-producing bacteria in brood cells and meconiums (first larval feces prior to pupation) of oil-collecting bees remains unknown. Here we provide new insight on the roles of these bacteria during diapause of an oil-collecting bee, *Centris aethyctera.* We also provide evidence that these antibiotic-producing bacteria may be within the entire guts of adult females (fore-, mid-, hindgut).

*Centris* spp. are bees nesting usually in the ground or in cavities (trees, dead branches, or human-made cavities). For ground-nesting species, the mother bee provides oil along with pollen to feed the larvae, cf. “oil-collecting bees”. The collected oil is used to adhere sawdust, sand, or soil to their legs, and serves to line the cells and seal the nest entrance [35]. When the cell is lined, the mother bee provisions the cell with a mixture of pollen, oil, soil material, and probably gland secretions at the bottom of the brood cell [36]. She then lays an egg on the provisions and seals the cell [35]. Nests of *Centris* spp. sometimes occur under extreme environmental conditions, i.e., high temperature, high humidity, and low oxygen. To protect the brood cells from parasites and pathogens, these bees need particular physiological and/or symbiotic adaptations [37]. Soils host a wide diversity of microbes and ground-nesting bees could benefit from gaining mutualistic microbes acquired from the environment. We hypothesized that bacteria capable of producing antibiotics, e.g. *Actinomycetes*, would occur mostly on/within the cocoon, where they may protect the pre-pupa against pathogens and parasites during diapause. To test for specific microbes in brood cells, we sequenced bacteria and fungi from multiple brood cell components (cocoon, meconium, and pre-pupa) of an oilcollecting bee, *Centris aethyctera*. We also tested for the presence of these bacteria in the gut of the *C. aethyctera* mother bees.

Our research contributes to the field of bee-associated microbiota at multiple levels. To our knowledge, this is also the first report of microbiotas (bacteria and fungi) of brood cell components of a solitary bee, *Centris* spp.

## Methodology

### Bee and brood cell collections and dissections

Bees and brood cells were collected using the permit #R-SINAC-SE-DT-PI-029-2023 (Costa Rica).

#### Brood cells of Centris aethyctera

On 24 April 2023 (1400-1600 GMTCST-6), 14 brood cells of *Centris aethyctera* (Fig. 1B) were collected from an open area within a mangrove swamp, with small patches of shrubs and herbaceous plants. *Centris aethyctera* commonly nested under *Sesuvium portulacastrum* and *Avicennia germinans*. The collection sites are near the town of Cuajiniquil, Guanacaste Province, Costa Rica (Bahía Tomas, site 1: 10°55’ 16” N, 85° 42’ 57” W, site 2: 10°55’ 15” N, 85° 42’ 56” W, Fig. 1A) and were formerly commercial salt evaporation ponds [38]. Both sites are characterized by a high degree of interannual variation (temperature and precipitation) and a dry season from December to May. During the highest tides, the regenerating mangrove experiences flooding during day and night, with some sites left covered in a layer of salt. Spotting the nest was only possible with the observations of Mauricio Fernández-Otárola a few months earlier when adults were still present, as evidence of nests disappears a few weeks after closure. Using sterile gloves, we manually excavated brood cells to a depth of 30 cm from the sediment where nests and flying adults were previously observed by Lobo et al. (2023), as evidence of nests disappears a few weeks after closure. Brood cells were placed in individual tubes, transported to the lab on ice, and then refrigerated at −20°C until dissection (about 2 days later). Each brood cell was dissected using sterilized tools under the stereomicroscope, and new collection samples were created to obtain: the cocoon, the meconium (i.e. first excreta of the larva before becoming pre-pupa into the cocoon), and the pre-pupa. *All* the collected brood cells were at the pre-pupal stage, except five brood cells with dead non-emerging adults (likely pharate adults from previous year); those were not considered in this study. Our collection time did not coincide with the presence of any adult *Centris aethyctera* at this location and date, April-May representing a *unique* phase of *C. aethyctera* at the pre-pupal stage. Additional information on nest and cell architecture of *Centris aethyctera* is in Lobo et al. (2023).

**Figure 1.**
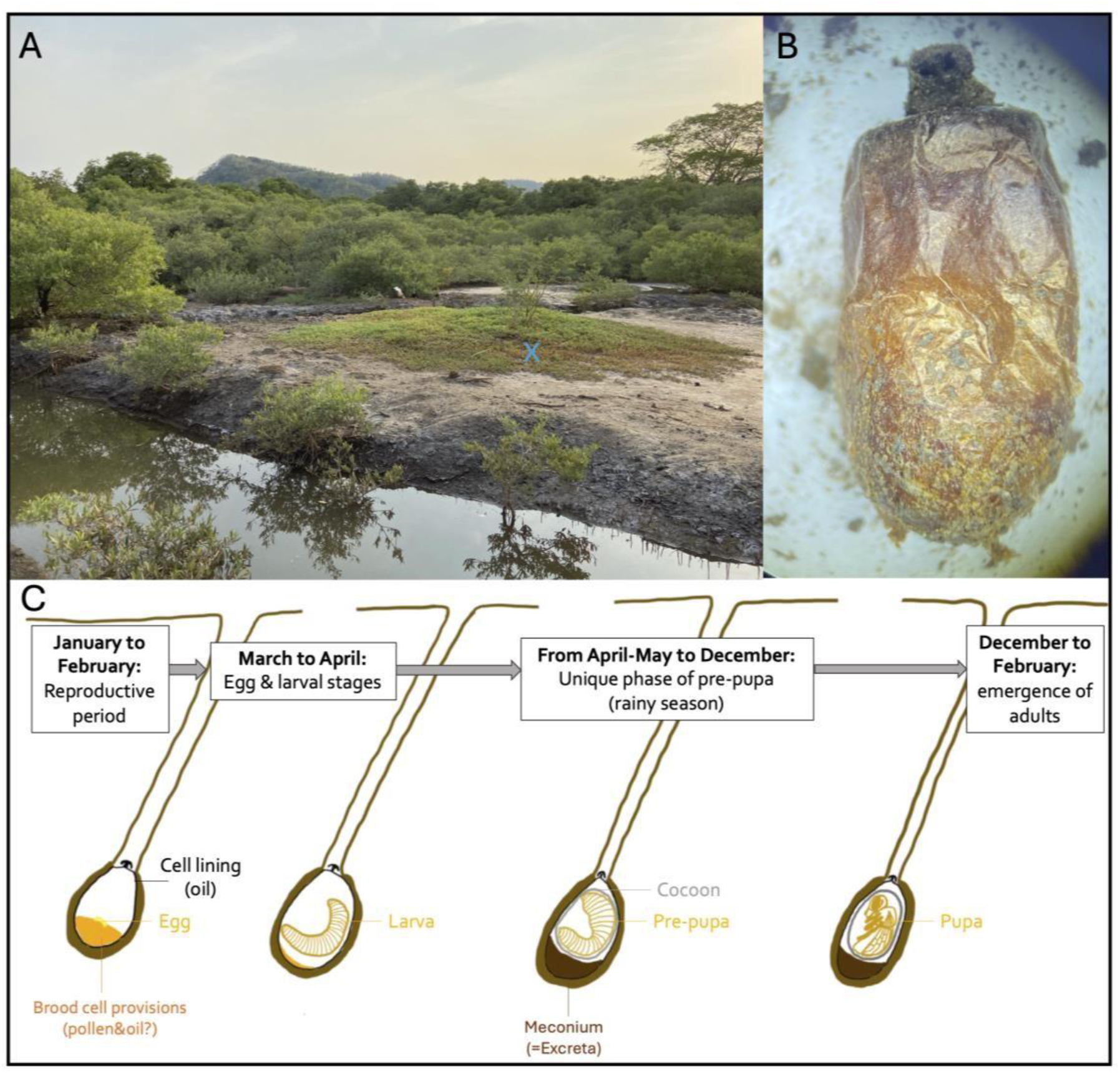
Panel A depicts *Centris aethyctera* nesting sites. The cross in yellow indicates the type of plant patches that cover the nest where the brood cells are located. The nest site is characterized by herbaceous plants fully covering the soil. In panel B, the image shows a Brood cell of Centris aethyctera detailing cell lining and the upper opercula for gas exchanges. Panel C the diagram shows the developmental stages of Centris aethyctera through the seasons: the reproduction period occurs from January to February when brood cells containing eggs and larvae are provisioned with pollen and probably oils; from March to April, larvae are more developed and have started to feed on the provisions and lastly, from April/May to December (rainy season), there is a unique phase of pre-pupa, where the individual is within the cocoon, after having digested the provisions and excreted it through the meconium (excreta). From December to February, the adults emerge, and males do so 2 weeks earlier than females.

#### Adult *Centris* spp. from Costa Rica

On 11 May 2023 (1500 1600 GMT-6CST), adults of three different species of *Centris* (*C. aethyctera* n=1, *C. flavofasciata* n=2, and *C. varia* n=9) were collected in an area covered with blooming and fruiting *Cojoba arborea* (wild tamarind) trees behind the beach of Playa Naranjo, in Guanacaste Province, Costa Rica (Bahía Nancite, 10°46’ 47” N, 85° 39’ 51” W). Bees were placed in individual tubes, transported to the lab on ice, and then refrigerated at −20°C until dissection (∼ 2 days later). The adult digestive tract (foregut to hindgut) of each individual was dissected using sterilized tools under the stereomicroscope. Only a single adult of *Centris aethyctera* was observed pollinating tamarind trees at that location and date, as adults of this species are usually absent during April-May. All samples from Costa Rica (brood cells and guts) were processed through cultureindependent methods (16S and ITS genes sequencing).

### DNA extraction & Microbiota analyses

#### DNA extraction

The DNA of all samples from Costa Rica (cocoon, meconium, and pre-pupa of *Centris aethyctera*, and guts of *C. aethyctera*, *C. flavofasciata*, and *C. varia*) were extracted using methodology described before [39]. The PowerSoil Pro Kit (QIAGEN LLC, Germantown Road, Maryland, United States) was used to extract DNA, following the manufacturer’s instructions, preceded by the addition of 20 µL of Proteinase K for 45 min at 60°C. The quantification of the extracted DNA was done using NanoDrop^TM^ 2000/2000c spectrophotometer (Thermo Fisher Scientific). The DNA concentrations varied for brood cells (average DNA yield=29.74 ng/µL) and adult guts (average DNA yield=136.42 ng/µL).

#### DNA amplification (16S rRNA and ITS genes)

The DNA obtained from all samples was then normalized to 4nM during the 16S rRNA and ITS genes library preparation. The Earth Microbiome Project standard protocols were employed for both bacteria and fungi (https://earthmicrobiome.org/protocols-and-standards/). The bacteria primers used were: 515F (5′GTGCCAGCMGCCGCGGTAA3′) and 806R (5′GGACTACHVGGGTWTCTAAT3′) to amplify the hypervariable region V4 of the 16S ribosomal RNA gene (∼291 bp) with regionspecific primers that include sequencer adapter sequences used in the Illumina flowcell. The fungal primers used were ITS1f-ITS2 (EMP.ITSkabir): forward primer (ITS1f) (5’-AATGATACGGCGACCACCGAGATCTACAC GG CTTGGTCATTTAGAGGAAGTAA3’) and reverse primer (ITS2), barcoded (5’-CAAGCAGAAGACGGCATACGAGAT NNNNNNNNNN CG GCTGCGTTCTTCATCGATGC-3’) to amplify the hypervariable ITS1 and ITS2 regions, that include sequencer adapter sequences used in the Illumina platform.

Amplified fungal regions varied in length (∼250-600 bp). Amplicons were quantified using PicoGreen (Invitrogen) and a plate reader (Infinite® 200 PRO, Tecan). Each product volume was then combined into a single tube to represent each amplicon in equimolar levels. AMPure XP Beads from Beckman Coulter were used to clean up this pool, and a fluorometer from Qubit from Invitrogen was used to quantify the results. Customized sequencing was outsourced to Argonne National Laboratory in Illinois, USA, utilizing a 2 x 250 bp paired-end sequencing kit with an Illumina MiSeq. The sequenced read and the associated metadata for brood cells of *Centris aethyctera* were uploaded to QIITA study ID 15303, and QIITA study ID 15451 for adults.

### Sequence processing and statistical analyses

#### Bacterial sequences

QIITA (version 2024.02) was used for initial processing of the resulting *fastq* files, i.e., demultiplexing, trimming at 250 bp, followed by denoising against the Greengenes2 database (version 2022.10) [40], using the deblur plugin (2021.09) in QIITA. Deblur method included quality filtering, removing chimeric sequences, combining pairedends, and eliminating singleton reads in order to join, denoise, and duplicate sequences. The obtained *.biom* files (without taxonomy) were processed locally in QIIME2 (version 2023.07) [41] and R (version 2023.11.1 build 402). The .*biom* files were first deblurred against SILVA (138 release, [42]), to obtain the DNA sequences of chloroplasts and mitochondria, currently not tagged in the 2022.10 Greengenes2 database. This subset containing chloroplast and mitochondria was then filtered manually form the initial .*biom* file using *antijoin* function from the *dplyr* package in R [43]. Singletons were also removed. The bacterial sequences were classified using the Greengenes reference database and taxonomy files [40], trained with classify-sklearn function in QIIME2.

#### Fungal sequences

Fungal sequences and their associated metadata were deposited in QIITA (version 2024.02) for repository, but the *fastq* sequences were processed in QIIME2 (version 2023.07). Sequences were demultiplexed and their quality score was assessed either considering paired-end sequences, i.e., forward and reverse, or single-end sequences, i.e., either forward or reverse. After paired-end vs. single-end quality score comparison, only forward sequences were chosen to proceed with downstream processes. DADA2 denoising method was used for quality filtering, chimeric sequence removal, single-end combinations, and singleton reads elimination to join, denoise, and duplicate sequences. The sequences were not trimmed, as fungal amplicons differ in base pair length. The fungal sequences were classified using the UNITE9.0 reference database and taxonomy files [44], trained with fit-classifier-naïve-bayes from QIIME2.

#### Meta-analysis of adult gut from *Centris* spp. from Costa Rica with adult *Centris decolorata* from Puerto Rico

Bacterial and fungal data sequences of *Centris decolorata* from Puerto Rico were retrieved from the QIITA study ID 14679 (retrieving only females, as only *Centris* spp. females were collected in Costa Rica), and then combined with the newly sequenced *Centris* spp. guts from Costa Rica into a new QIITA study 15451.

The downstream analyses were followed as in previous studies [45–47], with new analyses (functional and metabolic inference of bacteria and fungi). Microbiota analyses (diversity and composition) were divided into three sets: (1) brood cell samples of *Centris aethyctera* (cocoon, meconium, pre-pupa), (2) Actinomycetia from the brood cell samples of *Centris aethyctera* (cocoon, meconium, pre-pupa), and (3) guts of adult female *Centris* spp. from Costa Rica (*Centris aethyctera*, *C. flavofasciata*, and *C. varia*) and Puerto Rico (*C. decolorata*). For each set, beta- and alpha- diversity were calculated, and taxonomic compositions (taxa barplots) and putative biomarker taxa (MaAsLin and/or LEfSe) are represented. To correct for difference in extraction dates, those were added from the metadata as a reference for the MaAsLin analysis. In addition, for brood cells (set of analyses 1) and adults (set of analyses 2), core microbiome analyses and ecological and metabolic function inference (FAPROTAX for bacteria and FungalTraits for fungi) were computed.

#### Alpha and Beta diversity estimates

Alpha diversity analyses were computed using Faith’s Phylogenetic Diversity (PD) [48] and Shannon diversity index [49] as boxplots using the *microeco* package [50] in R (version 2023.12.1, build 402). Significant differences in richness and diversity were assessed using Kruskal and Wallis [51]. Beta diversity analyses were calculated using Bray-Curtis dissimilarity index and plotted using Non-Metric Multidimensional Scaling (NMDS) with samples colored according to the metadata categories. Different tests were applied to assess the significance of beta diversity measures: Permanova [52] and Anosim [53] to compare the dispersion of Bray-Curtis dissimilarity index in NonMetric Multidimensional Scaling; and Permdisp [54] to test the null hypothesis of homogeneity of multivariate variances.

#### Taxonomic compositions and putative biomarker taxa

Taxonomic compositions were illustrated using the *phyloseq* [55], *vegan* [56], and *ggplot2* [57] packages in R. Putative biomarkers differentially significant between multivariable associations were calculated using the *maaslin* package [58] in R. Another putative biomarker method was the Linear Discriminant Analysis (LDA) Effect Size (LEfSe), determining the features (at the genus level) most likely to explain differences between variables, by coupling standard tests for statistical significance with additional tests encoding biological consistency and effect relevance [59]. The parameters were alpha =0.05 (a significance level of 0.05 indicates a 5% risk of concluding that a difference exists when there is no actual difference) and logarithmic LDA score/threshold =2 (indicating the ranking of the biomarker relevance). This LEfSe analysis was computed and visualized using the *microeco* and *ggplot2* packages in R.

Terminology notes: The term *biomarker* will be used extensively in this microbiota study. We assumed its definition to be a characteristic microbial taxon (bacterial or fungal taxa) which is statistically measured (here with LEfSe and/or MaAsLin analyses) and assumed to be an indicator of the bacteria associated with a given class (here of each *Centris* spp. brood cell compartment and/or gut) [60–62].

#### Core microbiota

A core microbiota was identified for each variable (bacteria and fungi from brood cells and guts), using the *microeco* package in R. To evaluate the core microbiota, each taxon is displayed with heatmaps within the *microeco* R package according to its prevalence in 0 to 100% of the samples, and at different detection thresholds, i.e., the relative abundance of the taxon within the sample, compared to other taxa from the sample.

#### Correlation network analyses

Using SparCC method [63], a correlation network analysis of bacteria was computed including brood cell samples of *Centris aethyctera* (cocoon, meconium, and pre-pupa) in Microbiome Analyst 2.0 [64]. SparCC is a compositionally robust method, making a strong assumption of a sparse correlation network. It uses a log-ratio transformation and performs iterations to identify taxa pairs that are outliers to background correlations. Each node represents a taxon (at the genus level) and is illustrated through a pie-chart, indicating the prevalence of the concerned taxon in each sample, such that the larger the node, the higher the relative abundance of the represented taxon. Links represent the correlation of the taxon (node) with other taxa.

#### Inference of bacterial and fungal ecological and metabolic functions

Ecological and metabolic inference of bacterial functions, such as fermentation or nitrate reduction, were assigned to the bacterial genus with FAPROTAX 1.2.6 [65]. Ecological and metabolic functions with positive values only and those with a standard deviation higher than 0.01 were selected to compare the bacteria isolated from the brood cell samples of *Centris aethyctera*. For fungal function inferences, the FungalTraits database [66] was used within the *microeco* package in R, to retrieve information such as fungal guilds, fungal growth, or fungal environments. Functional annotations based on the FAPROTAX database only offer *inferred* functions based on 16S rRNA fragments. The precision of pathogen assignment in this study may not be as accurate as it would be in a true shotgun metagenomic study.

## Results

Given the arguments of McMurdie and Holmes [67], we decided to not rarefy the amplicon data (bacteria and fungi from brood cells or guts). Indeed, we expect in this study to unveil the presence of new bacterial taxa, especially from the Actinomycetia class which are sometimes not easy to identify using culture-independent methods [68]. Rarefying the samples would have increased the probability of missing these taxa from the different bee brood cells compartments and guts. In addition, in terms of microbiota, we were evaluating very different compartments within the brood cells. For instance, as pre-pupae excrete the content of their gut (meconium/excreta) before entering in dormancy within the cocoon, the microbial content tends to decrease in their guts. The read quantity for such sample is naturally lower than other samples from the brood cells (e.g., fungal reads from pre-pupae = 99,450.55 ± 54,487.75 versus 407,691.63 ± 155,093.30 in cocoons, **Supplementary Tables 1-2** and **Supplementary Tables 3-4**).

### *Centris aethyctera* brood cell diversity and composition

#### Alpha and beta diversity measures

The different compartments of *Centris aethyctera* brood cells (cocoon, meconium, pre-pupa) clearly show different alpha and beta diversities and composition. Microbiota (bacteria and fungi) from cocoon shows lower alpha-diversity using Shannon indices than that of meconium and pre-pupa (bacterial and fungal p < 0.05, **Figure 2 A-B**). While there is no difference in bacterial alpha-diversity between meconium and pre-pupa (Kruskal-Wallis test (K-W), p > 0.05, **Figure 2 A**), values for fungal alpha-diversity are significantly different, pre-pupae having a lower alpha-diversity in fungi (K-W, p < 0.01, **Figure 2 B**). When considering phylogenetic distances (shared branch lengths between bacteria), all brood cell components exhibit similar alpha-diversity, though meconium exhibits a greater intra-individual variation in phylogenetically different bacteria (Faith’s Phylogenetic Diversity index [PD], K-W, p > 0.05, **Figure 2 C**). However, fungal alpha-diversity analyses considering phylogenetic distances results in clear fungal compartmentalization between all brood cell samples, prepupae having the lowest value in phylogenetically different fungi, followed by cocoon (PD, KW, p< 0.05, **Figure 2 D**). Beta-diversity analyses disclosed similar pattern of compartmentalization between cocoon, meconium, and pre-pupa, for both bacteria and fungi (both permDISP p > 0.05, permANOVA, p < 0.001, anosim p-value<0.01, **Figure 2 E-F**).

**Figure 2.**
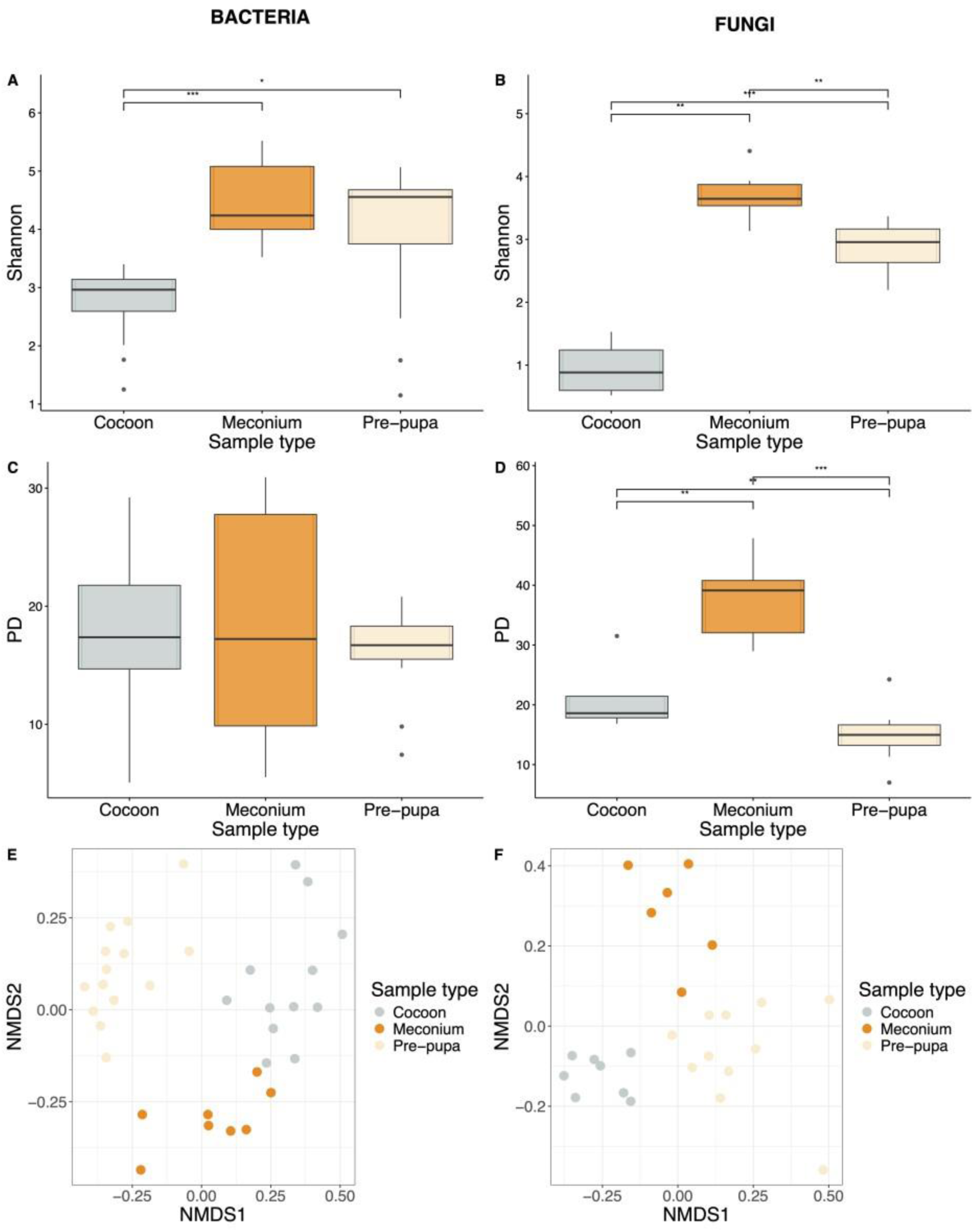
**Bacterial and fungal alpha and beta diversities of brood cell sample types of *Centris aethyctera***, i.e. Cocoon, Meconium, Pre-pupa: Bacterial alpha diversity analysis using Shannon index (A) and Faith’s Phylogenetic Diversity index –PD– (C), Fungal alpha diversity analysis using Shannon index (B) and using Faith’s Phylogenetic Diversity index –PD– (D), Bacterial (E) and fungal (F) beta diversity analyses using Bray-Curtis index. Stars indicate significance of Wilcoxon test: * indicates a p-value=0.05, ** indicates a p-value=0.01, and *** indicates a p-value=0.001.

### Taxonomic profiles and putative biomarker taxa

Actinobacteria, Bacteroidota, and Proteobacteria phyla are present in all brood cell samples (**Figure 3 A**), but Bacteroidota and Proteobacteria appear to be relatively more abundant in cocoon, Firmicutes in the meconium, and Campylobacterota in the pre-pupa (**Supplementary Figure 2 A**). The most abundant fungal phylum for all samples is Ascomycota (**Figure 3 C**), while Basidiomycota are present in the meconium and pre-pupa, and Mucoromycota occurs in the meconium (**Supplementary Figure 3 A**). When exploring finer taxonomic division, specific bacterial and fungal genera characterize different brood cell samples. Indeed, the LEfSe analysis demonstrated the presence of specific bacteria in each brood cell sample (alpha=0.05, min LDA score=3): *Marinobacter*, *Alcanivorax* and *Parvibaculum* characterized the cocoon; *Bacillus, Micromonospora* and *Clostridium* characterized the meconium; and *Escherichia*, *Pseudomonas* and *Chryseobacterium*, plus undescribed genera from the Enterobacteraceae family characterized the pre-pupa (**Figure 3 B, Supplementary Figure 1A**). These results were also confirmed with multivariate analyses (MaAsLin) applied for bacteria (**Supplementary Figure 2 B**). Fungal genera *Aspergillus Moleospora*, and *Septoriella* characterized the cocoon, *Arthrinium*, *Coprinellus*, and *Apiospora* characterized the meconium, and *Lecanicilium*, *Dothiora*, and *Aureobasidium* characterized the pre-pupa (LEfSe, **Figure 3 D;** MaAsLin, **Supplementary Figure 3 B**). The full lists of bacterial and fungal biomarkers are shown in **Figure 3 B & D** at the genus level, and in **Supplementary Figure 1 A-B** and **C-D** at the family and genus levels, respectively.

**Figure 3.**
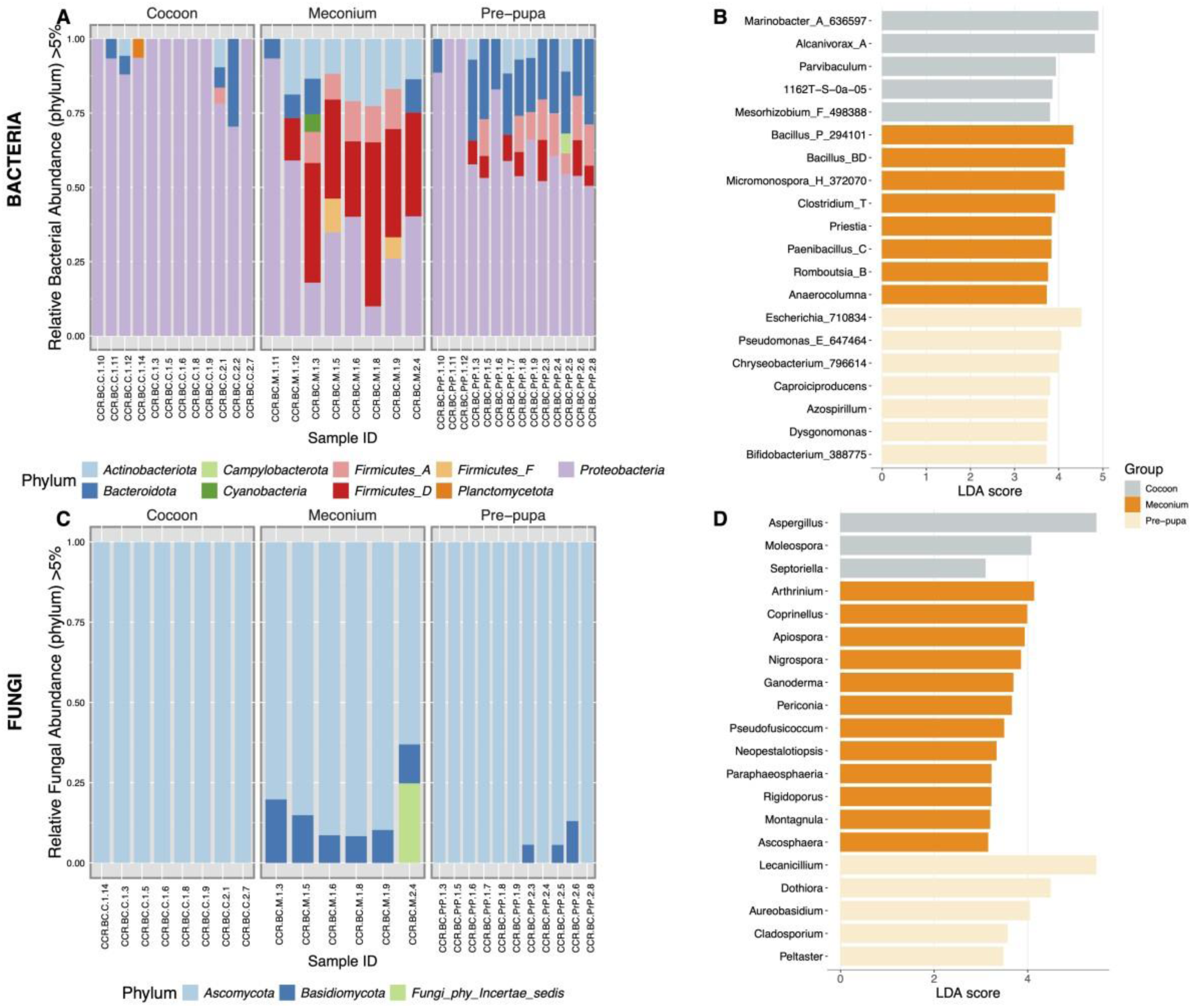
**Bacterial (A) and fungal (C) taxa composition of brood cell sample types of *Centris aethyctera***, i.e. Cocoon, Meconium, Pre-pupa, at the phylum level (showing all taxa having >5% of relative abundance); Linear discriminant analysis (LDA) effect size (LEfSe) at the species level of bacteria (B) and fungi (D) showing the top 20 significant taxa for each sample type (alpha=0.05, LDA score threshold=2). Undescribed genera are not shown in the LEfSe analysis.

### Core microbiota

Apart from these biomarkers, the core microbiota analysis reveals that each brood cell compartment has a set of bacteria and fungi genera that are present in 70-100% of all samples, each time representing up to 20% of the total sample abundance. Bacteriota taxa *Marinobacter* sp. and *Alcanivorax* sp. are present in at least 70% of all cocoon samples (> 1% relative abundance) (**Supplementary Figure 4 A**), *Bacillus* sp. and *Micromonospora* sp. are in at least 60% of all meconium samples (> 1% relative abundance) (**Supplementary Figure 4 C**), and *Escherichia* sp. (pre-pupal bacterial core) in 100% of pre-pupal samples at more than 1% of relative abundance (**Supplementary Figure 4 E**). In this core microbiota analysis, only the known genera are shown, the high abundance of unknown Enterobacteriaceae taxa in the prepupal samples is purposedly not shown.

In contrast to core bacterial biota, the core mycobiota within each brood cell sample type seems to be more obvious. *Aspergillus* sp. and *Leiothecium* sp. are present in 90% of all cocoon samples at min 10% of relative abundance (**Supplementary Figure 4 B**); *Lecanicillium* sp., *Leiothecium*, and *Dothiora* sp. are present in 100% of all meconium samples, at minimum 20%, 1% and 2.5% of relative abundance, respectively (**Supplementary Figure 4 F**); and *Lecanicillium* sp., *Leiothecium*, and *Dothiora* sp. are present in 100% of all pre-pupal samples, at minimum 20% and 2% of relative abundance, respectively (**Supplementary Figure 4 H**).

### Correlation network analyses

A network correlation analysis of bacteria from brood cells (SparCC method, p-value threshold = 0.01, correlation threshold = 0.7, **Supplementary Figure 5**) reveals that 1) bacteria from the cocoon has the highest bacterial abundance from all samples (**Supplementary Figure 2 A**), and 2) there is an apparent compartmentalization of correlated bacteria within the cocoon (**Supplementary Figure 5 C**), within the meconium (**Supplementary Figure 5 B**), and within the pre-pupa (**Supplementary Figure 5 C**) samples.

### Inference of bacterial and fungal ecological and metabolic functions

In addition to their bacterial compositional differences, brood cell samples also differ in ecological and metabolic functions. Ten predicted functional activities appear to be the most different between brood cell samples. Bacteria doing fermentation, nitrate reduction, photoautotrophy, and phototrophy are most abundant in the meconium and pre-pupae but are nearly absent from cocoons. Bacteria related to animal parasites and symbionts are observed in pre-pupae and meconiums. Cocoons have the highest bacterial function activity in (aerobic) chemoheterotrophy, aliphatic non-methane hydrocarbon degradation, and hydrocarbon degradation (**Supplementary Figure 6**). Observation of these patterns of ecological and metabolic functions in fungi are less obvious, while still exhibiting fungi characteristic of aquatic environments and fungi producing filamentous mycelium in all brood cell samples. Fungi from pre-pupae are characteristic of animal and (in-)vertebrate parasites or decomposers. In addition, fungal traits analysis report that these fungi initially originate from foliar endophytes (**Supplementary Figure 7**).

### Actinomycetia from *Centris aethyctera*’s brood cells

Actinomycetia, known to produce antimicrobial compounds, follows similar patterns of diversity compartmentalization. The alpha-diversity differences in Actinomycetia are the highest in the meconium, which has the highest richness (Chao1 index, K-W, p < 0.05) and diversity (Shannon index, K-W, p < 0.05) (**Figure 4 A**) compared to cocoon and pre-pupa samples. Beta-diversity analyses disclosed similar pattern of compartmentalization in Actinomycetia between cocoon, meconium, and pre-pupa (permDISP, p> 0.05; permANOVA, p < 0.001; anosim, p < 0.01, **Figure 4 B**). While cocoons are not characterized by any specific genus of Actinomycetia, the meconium is characterized by *Micromonospora*, *Actinomycetospora*, and unknown genera of Streptomycetaceae, and pre-pupa have *Bifidobacterium* (alpha=0.05, min LDA score=4, **Figure 4 C**). In addition, pre-pupal samples are composed mostly of *Cellulosimicrobium* (**Figure 4 D**). Based on culture-independent methods, actinomycetes seem to be prevalent in the meconium, and to a lesser extent in the pre-pupa, rather than in the cocoon. When computing a pattern search using Pearson r correlation index showing the top 25 bacteria, Actinomycetia from meconiums are indirectly correlated with the class of Cyanobacteria in the pre-pupa (**Supplementary Figure 8 A**). Using this pattern search for putative biomarkers of meconium, *Micromonospora*_H_372070 and *Actinospora* show a positive correlation with other bacteria from the meconium, namely with *Paenibacillus*_H*, Paenibacillus*_C, *Paenibacillus*_J_3668, and *Paenibacillus*_A (**Supplementary Figure 8 B-C**). *Bifidobacterium* spp. from pre-pupae are positively correlated with various other genera, such as *Janthinobacterium* spp., or *Lactobacillus* spp. *Bifidobacterium* spp. also show a positive correlation with genera present in the meconium, such as *Roseomonas* spp. (an acetic acid bacteria) or *Curtobacterium* spp. (**Supplementary Figure 8 D).**

**Figure 4.**
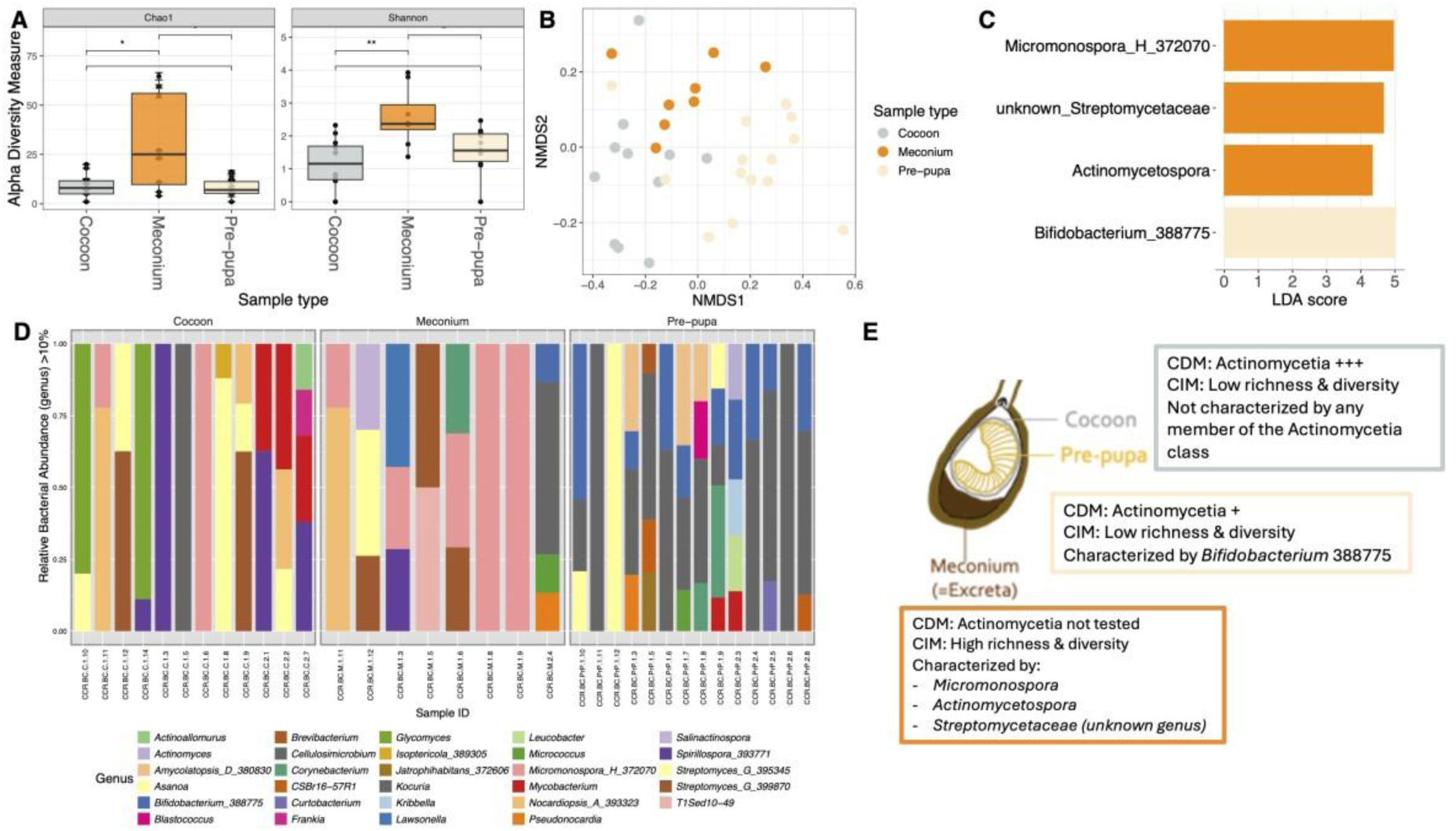
Diversity and composition analyses of Actinomycetia from brood cell samples of *Centris aethyctera*, i.e. Cocoon, Meconium, Pre-pupa. **(A)** Alpha diversity analyses using Chao1 and Shannon, stars indicate significance of Wilcoxon test: * indicates a p-value=0.05, ** indicates a p-value=0.01, and *** indicates a p-value=0.001; **(B)** Beta diversity analyses represented by a NMDS using Bray-Curtis index; **(C)** Linear discriminant analysis (LDA) effect size (LEfSe) at the genus level of the Actinomycetia class showing the top 30 significant taxa for each sample type (alpha=0.05, LDA score threshold=2). **(D)** Actinomycetia class composition at the genus level (showing all taxa having mora than 10% of relative abundance) for each brood cell sample types of *Centris aethyctera*, i.e. Cocoon, Meconium, Pre-pupa; **(E)** Summary of Actinomycetia compartmentalization within the brood cell samples of *Centris aethyctera*, i.e. Cocoon, Meconium, Pre-pupa, using culture dependent (CDM) and independent methods (CIM).

### Meta-analysis of gut microbiota from adult *Centris* of Costa Rica and Puerto Rico

The gut microbiota of all collected *Centris* (*C. decolorata* from Puerto Rico, and *C. aethyctera*, *C. flavofasciata*, and *C. varia* from Costa Rica) were sequenced, but because sample sizes were low for some species (**Supplementary Tables 3 & 4**), the diversity of the most densely sampled species are presented here, i.e., *C. decolorata* from Puerto Rico, and *C. varia* from Costa Rica. Alpha-diversity analyses of bacteria and fungi of adult female *Centris* guts resulted in a higher phylogenetic and Shannon diversities for the Costa Rican *Centris varia*, compared to *C. decolorata* from Puerto Rico (PD, K-W, p< 0.01, Shannon, K-W, p < 0.01, respectively, **Figure 5 A-D**). There is also a clear difference in beta-diversity between the two bee species for both bacteria and fungi (both permDISP, p > 0.05; permANOVA, p < 0.001, anosim, p < 0.01, **Figure 5 E-F**). Both species have a core bacteriome composed of acetic acid bacteria, particularly *Gluconobacter*. It is the most abundant bacterial genus, present in more than 80% of all *Centris* gut samples and representing up to 20% of the bacterial relative abundance per sample (**Figure 6 A & C**). This core bacteriome of acetic acid bacteria appears to be ubiquitous among all species of *Centris*, comprised of not only *Gluconobacter*, but also *Acinetobacter* and *Bombella* (**Supplementary Figure 9 A, C, E & G**). In contrast, our *Centris* samples revealed no clear core mycobiome (**Figure 6 B & D**). Nonetheless, *Aspergillus* and *Lecanicillium* are the most abundant fungal genera within *Centris* spp. of Costa Rica (*C. aethyctera*, *C. flavofasciata*, and *C. varia*), and are present in more than 90% of all samples, at a relative abundance up to 20% (**Supplementary Figure 9 D, F & H**). In *Centris aethyctera*and *C. flavofasciata*the core gut mycobiome includes undescribed species of *Penicillium*, which is present in 100% of the guts at low relative abundance (up to 4.5%, **Supplementary Figure 9 F & H**).

**Figure 5.**
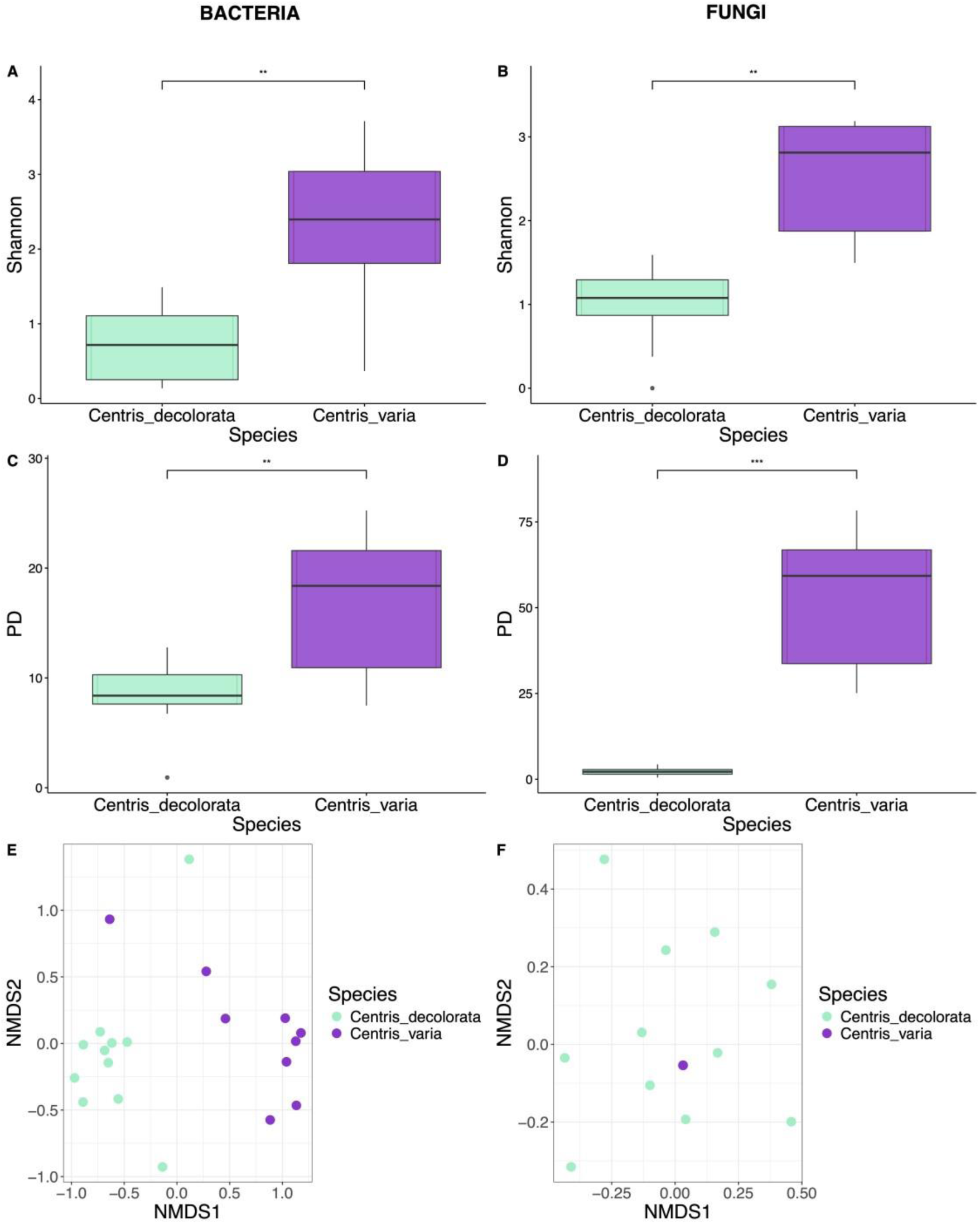
Bacterial and fungal alpha and beta diversities among adult individuals *Centris decolorata* (Puerto Rico) vs. *Centris varia* (Costa Rica) female guts: Bacterial alpha diversity analysis using Shannon index (A) and Faith’s Phylogenetic Diversity index –PD– (C), Fungal alpha diversity analysis using Shannon index (B) and using Faith’s Phylogenetic Diversity index –PD– (D), Bacterial (E) and fungal (F) beta diversity analyses using Bray-Curtis index. Stars indicate significance of Wilcoxon test: * indicates a p-value=0.05, ** indicates a pvalue=0.01, and *** indicates a p-value=0.001.

**Figure 6.**
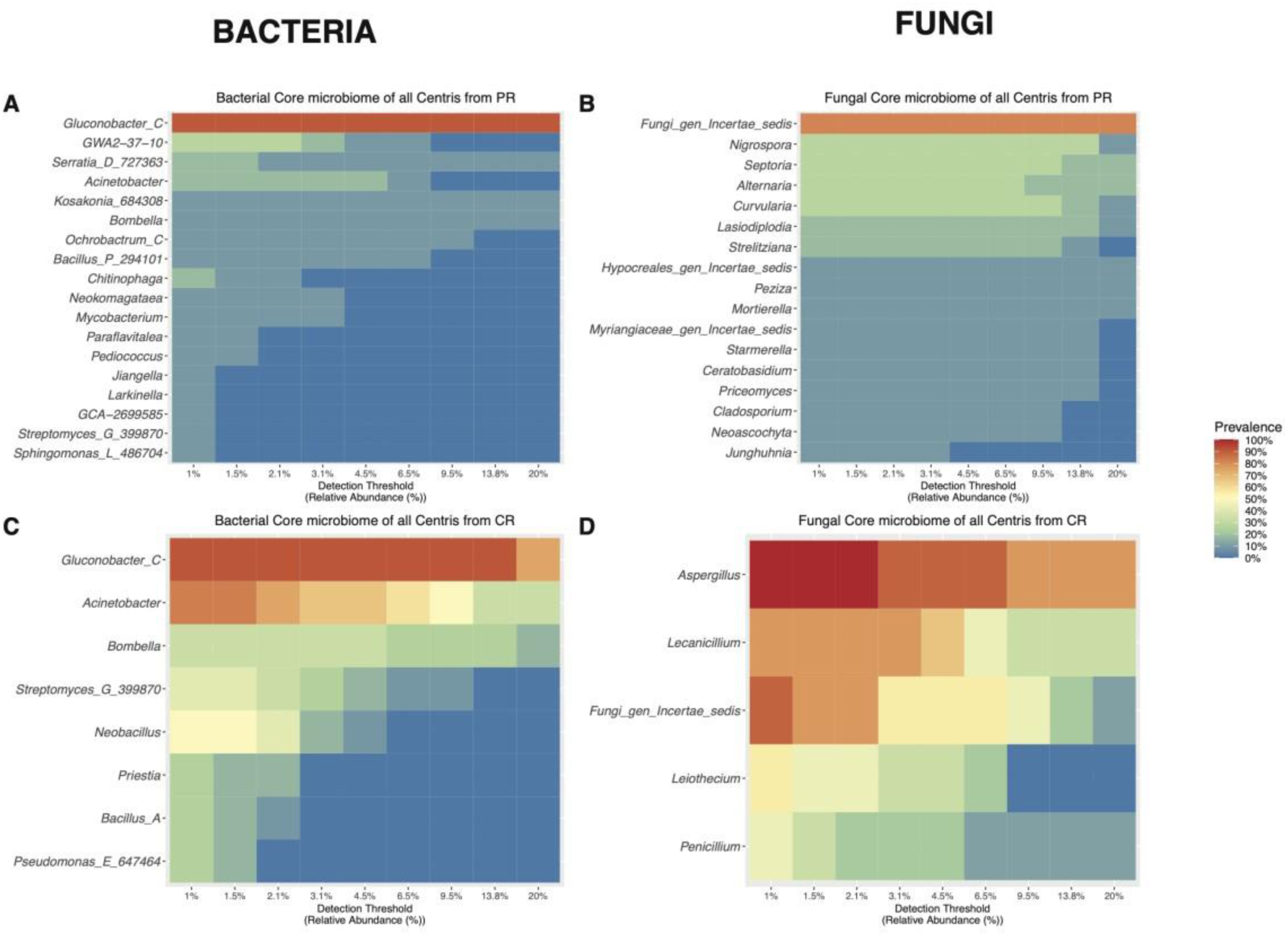
**Heatmaps of Core Microbiome among female *Centris* guts from Costa Rica vs. Puerto Rico,** at the genus level (y axis). Colors indicates the prevalence of the taxa in the samples, from blue/0% (present in 0% of all the samples) to red/100% (present in 100% of the samples), with a certain detection threshold (x axis) indicating the relative prevalence in % of the taxa within the sample. Panels A and B represents the heatmaps of bacterial core microbiome in all *Centris* from Puerto Rico, while Panels C and D represent the heatmaps of fungal core microbiome from all Centris from Costa Rica.

## Discussion

Microbiota of bees and their nests can have critical functions that facilitate and/or ensure bee survival through antimicrobial effects on deleterious microbes. While characterization of microbiota of social bees is advanced, that of solitary bees remains largely unknown. In bee nests, the transmission and compartmentalization of these microbes may be regulated by the activities of the mother bee. Knowing the diversity and natural history of such beneficial microbes may inform strategies to protect these critical components of ecosystem sustainability, but may also generate a new class of antimicrobial agents.

We investigated the presence of bacteria and fungi in the different brood cell compartments (cocoon, meconium, pre-pupa) during pre-pupal diapause, as well as in the mother bee gut of multiple species of *Centris*, a genus of oil-collecting bees. We found that 1) there is a clear difference in diversity and compositions of both bacteria and fungi between the different brood cell compartments, 2) brood cell compartments possess antibiotic-producing microbes, whether actinomycetes or not, 3) the meconium (pre-pupal fecal matter) contains most of the antimicrobial-producing microbes (rejection of our initial hypothesis of prevalence of actinomycete bacteria on/within the cocoon layers), and 4) *Centris* females (*i.e*., mother bee) contain in their guts some potentially beneficial microbes also present in the brood cells.

Because our results come from 16S and ITS sequencing, we can **only speculate on the possible bacterial and fungal functions** in each brood cell compartment and mother bee gut. Our following inferences on the potential significance of these microbe functions are thus **purely hypothetical** and future research should specifically test the actual functions of these microbes.

### Microbial defense within the brood cells Bacteria-owed defense

#### Cocoon

Most bacteria living on or within the cocoon layers are characteristic of oceanic environments, especially taxa usually found in oil-contaminated environments, e.g., *Alcanivorax dieselolei* [69] or *Oleiagrimones citrea* [70] that can use hydrocarbons as a carbon source in the presence of oxygen [69,70]. The discovery of such microbes in a recovering mangrove swamp is not surprising, after so many years of anthropogenic stress (salt extraction site). However, the selection of these bacteria in the cocoon compartment might appear quite surprising. However, at this mangrove site, oxygen penetrates the substrate to less than 1 mm, yet the brood cells were extracted from the substrate 5-13 cm deep [71].

Oxygen is even less available when floods occur on a recurrent basis [71]. Both bees and their heterotrophic microbes, such as these alkane-degrading bacteria, i.e. *Alcanivorax dieselolei* or *Oleiagrimones citrea*, require oxygen to survive. Hydrocarbon degradation under anaerobic conditions is possible in the presence of oxygen providers, such as nitrogen oxides (NO_x_) [73]. In fact, we found that the cocoon contains bacteria involved in the nitrogen cycle: nitrogen fixing bacteria (N_2_ to NH_3_) such as *Mesorhizobium* spp. [74] and nitrate reducing bacteria (NO_3_ to NO_2_) such as *Solirhodobacter olei* [75] or *Pelagibius litoralis* [76]. As a source of carbon, these bacteria may use glucose from the degradation of cellulose (issued from previous pollen intine digestion, i.e. internal layer of the pollen wall) by Cellvibrionaceae bacteria [77], also constituting a biomarker of the cocoon (*cf*. “1162T-S-0a-05” genus from the Cellvibrionaceae in the cocoon). We hypothesize that these metabolic reactions coupling hydrocarbon degradation and nitrogen fixation/reduction could be essential for the survival of bacteria such *Alcanivorax dieselolei.* As a product of hydrocarbon degradation, *Alcanivorax dieselolei* produces a biosurfactant [78], which could simply act as a physical barrier against desiccation of the cocoon and the pre-pupae but could also interfere with the production of other bacterial biofilm [79], reducing the probability of pathogenic infection of the pre-pupa.

Another important biomarker and core microbiota constituent of the cocoon is *Marinobacter bohaiensis*, which is associated with the production of hydrogen sulfide. This molecule could have several important chemical implications, including providing antibiotic resistance [80]. This property may allow *M*. *bohaiensis* to exist in proximity to antibiotic producing bacteria of the meconium. Gas-mediated antibiotic resistance could be shared with other bacterial members of the cocoon. Thus, some bacteria from the cocoon could synergistically interact to provide physical and chemical protection to the pre-pupa.

#### Meconium

Given culture independent approaches, meconium is the brood cell compartment with the highest diversity and abundance in actinomycetes. *Micromonospora* spp., a biomarker of *Centris aethyctera*’s meconium, can produce a variety of antimicrobials, such as aminoglycoside, enediyne, and oligosaccharide antibiotics [81] but also other active molecules such as vitamin B12 [82] and anti-fungal compounds [83]. While not reported from either pollen or meconium before, *Micromonospora* has been previously isolated from nursing stingless bees (*Melipona scutellaris*). These bacteria, along with *Streptomyces* spp., were shown to have the ability to inhibit the growth of *Paenibacillus larvae*, the causative agent of the American Foulbrood disease in honeybees [84]. Although *P. larvae* was only present in trace amounts in *Centris aethyctera* brood cells, other *Paenibacillus* taxa were present, including undescribed and described species (such as *P. populi*, for which antimicrobial properties have not yet been described [85]). *Paenibacillus* has been widely described from wild bee surface bodies and guts [86], and especially within nests of megachilids [86–88], without showing any negative effects in brood cells. In fact, some strains have been shown to have chitin-binding, biofilm-forming, and antimicrobial capabilities [86,89–91]. These characteristics could be useful to avoid fungal penetration of the bee cuticula, and the presence of *Paenibacillus* in bee nests could be beneficial to reduce fungal threats in humid and nutrientrich environments of wild bees [86], such as *Centris* spp. However, better taxon cultivation and genomic description are required to infer beneficial effects of *Paenibacillus* bacteria within the brood cells of *Centris*. Other meconium biomarkers include *Bacillus*_P and *Bacillus*_BD, taxa that have been shown to produce a wide variety of antibiotics [92], especially against phytopathogenic fungi [93]. Yet again, full species description of the *Bacillus* genus should be done to confirm the capability of these bacteria to produce antimicrobials and to know their precise targets.

In *Centris aethyctera* meconiums, Actinomycetes could control the presence of Cyanobacteria in the brood cells. Photosynthetic cyanobacteria are mostly present in aquatic environments, and more than half can produce toxins. This inhibition appears balanced, rather than complete, as cyanobacteria are still present in the pre-pupa. Cyanobacteria have been observed previously in healthy bee guts, e.g., queen bumblebee [94], honeybee [95], and their toxin does not impact larval mass or adult survival in honeybees [96]. In our study, the presence of cyanobacteria is low and represents less than 5% of relative abundance in pre-pupae. Furthermore, most of them are undescribed genera and species, and their level of toxicity is thus not known.

#### Pre-pupa

Describing the potential microbial defense properties provided by bacteria in *Centris aethyctera* pre-pupa is quite difficult. Most bacterial taxa are uncultured or classified up to the family level only. Bacteria from the Enterobacteriaceae and *Pseudomonas* spp. have been already observed in digestive tracts and nests of bees, e.g., [8,34,37,97]. *Chryseobacterium* spp., another bacterial genus constituting the pre-pupal core microbiota, are often associated with a multitude of hosts, including insects, e.g., [98]; but to our knowledge, it is the first time it has been isolated from bees. Known species from this genus could inhibit plant pathogenic fungi [99], degrade nematodes and other helminths [100], and even produce toxins such as botulinum neurotoxins, as some homolog genes are present in their genomes [101]. This genus also shows high resistance against antibiotics [102], which could provide an advantage given the presence of antibiotic-producing *Bifidobacterium* spp. in the pre-pupa. This genus is an actinomycete biomarker of *Centris aethyctera* pre-pupae, and some strains produce a variety of antimicrobials that have immunomodulatory effects [103]. It can degrade biofilms produced by a multi-drug resistant *Escherichia coli* [104], or even inhibit the growth of *Clostridioides difficile* in combination with other antibiotics [105]. It is interesting to note that *Clostridioides difficile* was found in *Centris aethyctera* meconium, but not in the pre-pupa.

### Fungal-owed defense in the brood cell

The role of fungi in protecting *Centris aethyctera* brood cells is quite challenging to ascertain. First, metabolic functions of microscopic fungi are largely unknown compared to macroscopic fungi, and most studies generally concern pathogenic ones. Second, interest in non-pathogenic fungi of bee microbiotas is relatively recent, but there is some indication that fungi may benefit bee larval development. In the stingless bee *Scaptotrigona depilis*, *Zygosaccharomyces* fungi are important for larval survival, as the fungi produce ergosterols, a precursor of hormone biosynthesis, necessary for pupation [107,108]. While a recent review on bee fungal associations studies failed to reveal any instances of fungi providing protection to bees, the data are sparse, and no study addressed bee-fungal interactions of solitary bees [109].

In *Centris aethyctera* brood cells, fungal presence is reduced, in contrast to other tropical bees, e.g., *Ptiloglossa* [21]. Because of the absence of lactobacilli proposed to be controlling fungal growth in the tropical bee *Ptiloglossa*, *C. aethyctera* brood cells display a high fungal diversity and composition, specific to the different compartments of the brood.

#### Cocoon

Fungi are predominant in the cocoon, namely *Aspergillus* spp. Some, especially *A. niger and A. japonicus* [110], can produce citric acid, which can be a carbon source used by *Marinobacter bohaiensis* [111] and *Alcanivorax dieselolei* [69], bacteria also present in the cocoon. Another one, *Aspergillus melleus*, produces neoaspergillic acid as well as mellein, compounds that are known to be antifungal [112]. Neoaspergillic acid is also known to be antibacterial and has been previously isolated from marine-derived mangrove epiphytic fungi [113], as well as mullein. These species of fungi have potent antifungal effects against *Alternaria alternata*, *Botrytis cinerea*, and *Fusarium oxysporum,* some opportunistic phytopathogens [112]. Other *Aspergillus* fungi could control ambient acidity by using their organic acid secretions [110]. For instance, *A. japonicus* has been shown to inhibit phytopathogenic fungi *Sclerotinia sclerotiorum* by releasing acid compounds [110]. The presence of these *Aspergillus* species might thus represent an advantage by providing metabolites for bacteria (e.g., citric acid), producing antifungal compounds against other phytopathogenic fungi, and controlling the ambient acidity.

#### Meconium

Fungi associated to the meconium originates naturally from plants. Most of the meconium biomarkers are endophytic fungi, such as *Coprinellus* spp. and *Apiospora* spp. More specifically, *Coprinellus magnoliae* is an endophytic fungus isolated from *Magnolia candolli* [114]. The prevalence of fungus from the Basidiomycota phylum highlights their potential to degrade cellulose-rich substrates [115], such as residual pollen present in the meconium. The product of cellulose degradation by these fungi is cellobiose [116], which could also be used as a source of carbon by other bacteria, namely *Nialli oryzisoli* [117], also present in the meconium as a biomarker. *Arthrinium phaeospermum*, another important biomarker from the meconium, is known to be a globally distributed pathogenic fungus with a wide host range, i.e., plants, humans, and animals [118]. However, some strains producing gibberellins could promote plant growth [119]. These results show the importance of assessing the pathogenicity of microorganisms, especially fungi, at the strain level, as it has been suggested before [109].

#### Pre-pupa

The presence and dominance of *Lecanicillium coprophilum* in the pre-pupae and the mother bee gut could be an argument in favor of the role of symbionts in larva development. Indeed, *Lecanicillium coprophilum*, a fungus isolated for the first time from feces of a marmot [120], can prevent and control diseases induced by *Cordyceps militaris*, an entomopathogenic fungus [121]. This latter zombie fungus attacks insect larvae and suppresses their immune response [122,123]. Because the transmission mode of *C. militaris* depends on killing its host [122], its population control and prevention in *Centris aethyctera* brood cells might be essential. While it is tempting to infer the beneficial role of *Lecanicillium coprophilum* presence in brood cells and guts of *Centris* spp. from Costa Rica, further research should investigate the precise mechanisms by which *Lecanicillium coprophilum* would benefit from the symbiose.

### Fermentation as a defense tool in brood cells of *Centris aethyctera*

Indirectly, bacterial fermentation occurring in the guts and nests of wild bees [124,125] could also suppress microbial invaders. Indeed, their products are often acidic, either due to lactic acid bacteria (e.g., [21]) or acetic acid bacteria (e.g., [34]). In *Centris aethyctera* brood cells, the highest fermentation activity by bacteria is observed in the pre-pupa, followed by the meconium. The cocoon does indeed contain fermenting bacteria, such as *Bifidobacterium* spp. or *Dysgonomonas algaminatilyca;* and the meconium also has fermenting bacteria, e.g., *Anaerocolumna, Niallia oryzisoli*, or *Hydrogenispora ethanolica*. Some of the subproducts of these bacteria are acids [126–128], which have the potential to increase the acidity in the brood cell, decreasing the survival potential for microbes unable to grow in acid rich environments. Thereby, these acidic subproducts considerably decrease the pH of the environment, acting as a pathogen filter.

### Interaction between fungi and bacteria in *Centris aethyctera* brood cells

We did not directly test the interaction between fungi and bacteria through metabolic pathway analyses. Nonetheless, the extensive literature review on the respective biochemical activity of bacterial and fungal biomarkers from each sample exemplified the processes by which bacteria and fungi could interact within the brood cells. This literature review had also the merit to direct future hypothesis testing. In particular, future experimental designs involving indiapause brood cells should manipulate the presence of specific biomarkers, such as the potential of *Lecanicillium copronilus* in preventing cordyceps disease in *Centris aethyctera* larvae, or the suppression of actinomycete bacteria to study the extent by which they confer antimicrobial protection to the brood cells. Future investigation should also consider the biochemical composition of each brood cell compartment. To our knowledge, only one study analyzed the chemical composition of a solitary bee cocoon, a megachilid [129]. The importance of substrate chemical composition would thereby be directly linked with the metabolism of each bacterium and fungus, giving importance to the intercorrelation between microbes and/or with the host material.

### Good parenting from *Centris* mother bees: symbiont provisioning?

Because they are holometabolous insects, *Centris* bees completely change their morphology between life stages, i.e., egg, larva, pre-pupa, pupa, pharate and adult. Prior to diapause and pupation, larvae defecate and purge their gut, along with its microbes. These include microbes that helped with the larval digestion of brood cell provisions, those that were potential pathogens, and those that protected the larva from pathogens and parasites. This microbial reduction is an advantage in the prevention of infections by pathogenic microbes, but a disadvantage by reducing or eliminating beneficial symbionts [130]. Trade-offs likely occur between limiting the spread of parasites and pathogens and creating a friendlier environment for symbiont persistence [131]. For instance, some lepidopteran hosts and their symbionts are able to interact to allow the passage of symbiotic bacteria through pupation, and to avoid the opportunistic infection during the larva-pupa metamorphosis, by up-regulating its immune function [131]. Other holometabolous insects overcome this potential loss of symbionts by having overlapping generations within nests, where they can transfer the symbionts between individuals of the nest, such as it is the case for eusocial hymenopterans, e.g., *Acryomyrmex* leafcutter ants, honeybees, or bumblebees [132–135]. In solitary hymenopterans such as beewolf wasps, larvae can integrate symbionts from the brood cell walls to the cocoon, that the metamorphosed adult would intake before emerging [136]. While mechanisms of maintenance of symbionts during the passage between larvae and pupae in solitary bees remain to be studied, similar strategies could occur in *Centris* bees. The persistence of beneficial symbionts in adult stage may be allowed by the persistence of the symbionts in non-gut organs/structures of the pre-pupal body, followed by its migration after the structure disintegration during metamorphosis, as observed in other holometabolous insects [137]. Some of the symbionts could also escape the gut before defecation, to be reinoculated later [138]. Most of the time, a minute inoculum is often sufficient to allow the symbiont passage from larvae to pupae [139], and beneficial bacteria might be selected and retained to the next life stage.

In that sense, the microbes coming from the mass provisioning could be the ones mediating metamorphosis within the brood cell by protecting the pre-pupae against pathogens. Since *Centris* mother bees lay their egg at the surface of their mass provisioning, the symbiont transmission *could*, to some extent of the definition, be considered vertical. The egg is literally bathed in a pool of potential symbionts, which would be ingested after developing into a larva. Once the entirety of the mass provisioning is consumed, the larva will defecate the digested content, along with the inhabiting microbes, just before the diapause, and form a cocoon in which it will perdure for 9 months. The mass provisioning could contain beneficial microbes, protecting the pre-pupa during the vulnerable diapause time.

Because the meconium of this present study contains a high abundance of antimicrobialproducing microbes, it could represent a first barrier to avoid the spread of parasites and pathogens into the metamorphosing pre-pupal gut. While the mother-provisioned symbionts are absent or greatly decreased from within the pre-pupa and pupal bodies, we show here that there might be a selection for bacterial and fungal symbionts having the capacity to protect the spread of pathogens to the bee (pre-pupa, pupa, pharate). Second, the cocoon also contains antibiotic-producing actinomycetes, according to preliminary results of culture-dependent methods, which could represent a second barrier of protection against pathogen migration. And finally, the pre-pupa itself still contains antimicrobial-producing microbes, protecting itself against pathogenic establishment in its tract.

However, this vertical transmission of symbionts may only serve as a *starting pack of symbionts*, which would be greatly reduced or lost entirely following moulting. Indeed, the complete metamorphosis and the divergence of life stages (egg, larva, pupa, pre-pupa, and pharate within soil/cavities vs. complete adults pollinating) imply the difficulty in maintaining vertical transmission of symbionts in *Centris*. Rather than strictly using vertical transmission routes, *Centris* are more likely to use alternate strategies for inter-generational transmission, and/or to acquire symbionts from the environment. For instance, the presence in the brood cell *and* the mother bee gut of Streptomycetaceae bacteria could be simply a result of horizontal transmission routes. The pre-pupa, once becoming an adult bee, may simply reacquire these potential symbionts by visiting the same floral resources as other adults from the same species, but it could also re-inoculate itself before emerging, as it has been shown for beewolf wasps.

### Fitness advantage for microbes despite life stage bottlenecks

In our study, symbionts are probably not issued from a co-evolution with their hosts. Indeed, for solitary bees, the strict co-evolved host/symbionts would be disadvantageous [130]. Here most of the bacteria present in the brood cells and guts originate from the soil. The arrival of rich substrate, such as pollen, pre-digested pollen, aerial plant material, could present an advantage for the microbes of the soil, in addition to the host. Even though the host undergoes metamorphosis, their dispersal is not strictly necessary as they could simply stay in the soil, maybe with a lower fitness given the lesser substrates in absence of nests. Due to the limitation of our amplicon sequencing study, it remains to be tested whether there might be a recycling of these symbionts across bee generations providing an advantage for these symbionts.

### Extreme environment ground-nesting bee nests as a source for novel antibiotics discovery

Antibiotic resistance affects 2 million people in the US, requiring $20 billion in excess healthcare costs (Center for Disease Control and Prevention, CDC) making it an important public health crisis. Whether the resistance is towards a single or several antibiotics, resistance occurs due to selective pressure on bacterial populations. New antimicrobials are needed to counter resistance and yet no new antibiotic classes have been clinically approved in over three decades. Over the last 80 years, *Streptomyces* spp. have been a good source for novel antibiotic compounds. Indeed, it represents the most economically and clinically important bacterial genus. They produce the most important antibiotics, including tetracycline, streptomycin, or erythromycin. Among the wide variety of secondary metabolites they produce, two-thirds represent the commercially available antibiotics. But since 1985, soil has been the single source of discovery of only three new families of *Streptomyces*-derived antibiotics. In the light of antibiotic resistance rise, developing new medicines is critical. To respond to the call from the World Health Organization (WHO) for the development of novel antibiotics to treat drugresistant infections, it becomes necessary to screen *Streptomyces*-derived antibiotics in environments other than soils. Natural metabolites and molecules have evolved to moderate communication between microbial species. The diversity of natural product chemistry seen in microbiomes has emerged a new paradigm in the identification of antimicrobial drugs within this host-microbe interactions. Insect-associated *Streptomyces* spp. have already been proposed as a new source of antimicrobials. The diversity of these compounds is expected to be high; since millions of years of evolution, insects have been bioprospecting continuously these active and defensive molecules, as a result of the selective pressure for pathogen inhibition. For humans, that may represent an opportunity to develop new medicines, as these compounds are also expected to have low toxicity to animals, as they are to other eukaryotic cells [18]. Here, we propose more specifically brood cells from ground-nesting bees as a new source of antibiotics. We demonstrated a new environment, rich in actinomycetes, as well as other antimicrobial producing microbes. These microbes deserve better attention, as most of them are undescribed families, genus, species, and strains. Their description and cultivation could lead to the exploration of totally new classes of antimicrobials, necessary to answer the call for new medicine discovery by the WHO.

## Conclusion

Given the results of culture-independent methods, we show that the brood cells of oil-collecting *Centris* bees represent an antimicrobial rich environment. In addition to a possible microbial migration from the soil, this special microcosm is probably engineered by the mother bee through mass provision. Indeed, some of these microbes (namely Streptomycetaceae and *Lecanicillium copronillus*) are also present in the mother bee gut. The non-random distribution of microbes across the different brood cell components suggests that they may have specific functions. While our study relies solely on inductive reasoning, with inference of the respective microbial metabolic activities, this first characterization of an oil-collecting bee brood cell certainly leads to new hypotheses. Further research is necessary since these microbiological conditions are probably crucial for pupal development in extreme environments. We believe that more research on the microbial interaction networks in brood cells of solitary bees will enhance our knowledge of the variables influencing microbe population dynamics and enable us to investigate fitness components that have not yet been identified.

## Supplementary Tables and Figures

**Supplementary Table 1.** Summary of variables for the brood cells of *Centris aethyctera* (Cocoon, Meconium, Pre-pupa) from Costa Rica considering bacteria.

| Brood cell<br>*Centris*<br>*aethyctera* | Sample<br>type | Nb of<br>sample | Avg.<br>read/sample | Read st. dev. | Avg.<br>OTU/sample | OTU st. dev. |
| --- | --- | --- | --- | --- | --- | --- |
|  | Cocoon | 12 | 10737.25 | 4880.45317 | 207.9166667 | 84.2576766 |
|  | Meconium | 8 | 1470.875 | 1115.7225 | 247.25 | 181.800322 |
|  | Pre-pupa | 14 | 1635 | 558.320421 | 158.5 | 53.6351636 |
|  | **TOTAL** | **34** |  |  |  |  |

**Supplementary Table 2.**
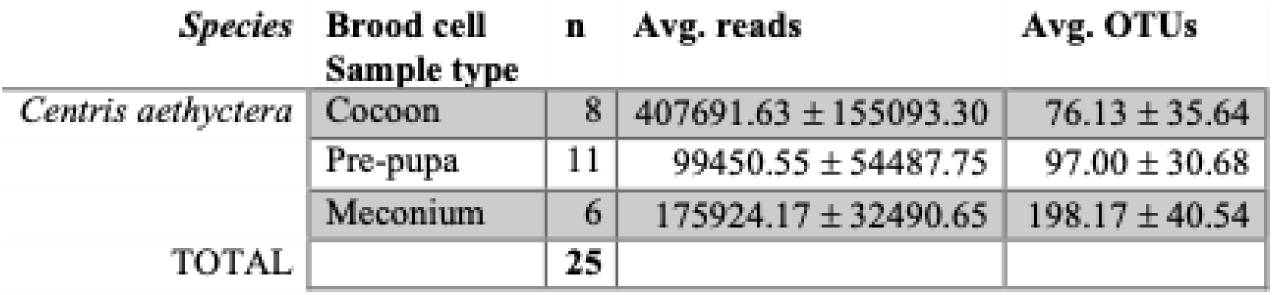
Summary of variables for the brood cells of *Centris aethyctera* (Cocoon, Meconium, Pre-pupa) from Costa Rica considering fungi.

**Supplementary Table 3.** Summary of variables for the different species of female *Centris* (*C. decolorata* from Puerto Rico; *C. aethyctera, C. varia,* and *C. flavofasciata* from Costa Rica) considering gut bacteria. Species with individual number n≤2 (*Centris aethyctera* and *C. flavofasciata*) are considered for taxa composition display (taxa barplots of phylum and genus) but are retrieved for further analyses (alpha and beta diversity analyses, MaAsLin, and FAPROTAX).

| Location | *Species* | n | Average reads | Average<br>features |
| --- | --- | --- | --- | --- |
| PR | *Centris decolorata* | 11 | 2710.09 ± 3193.08 | 22.70 ± 14.14 |
| CR | *Centris aethyctera* | 1 | 1907.00 ± 0.00 | 51.00 ± 0.00 |
|  | *Centris varia* | 9 | 4596.00 ± 3208.35 | 125.44 ± 60.66 |
|  | *Centris flavofasciata* | 2 | 10701.00 ± 11803.03 | 57.00 ± 49.50 |
|  | TOTAL | 23 |  |  |

**Supplementary Table 4.**
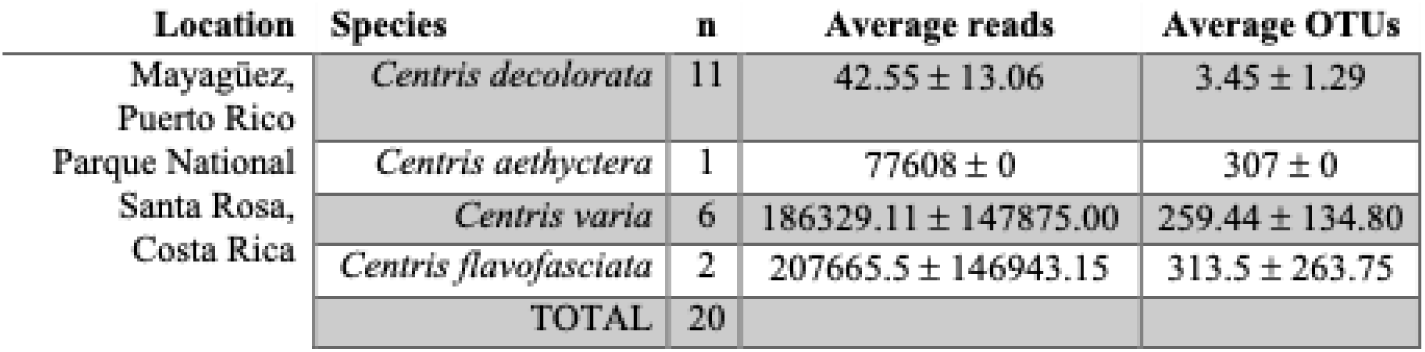
Summary of variables for the different species of female *Centris* (*C. decolorata* from Puerto Rico; *C. aethyctera, C. varia,* and *C. flavofasciata* from Costa Rica) considering gut fungi. Species with individual number n≤2 (Centris aethyctera and C. flavofasciata) are considered for taxa composition display (taxa barplots of phylum and genus) but are retrieved for further analyses (alpha and beta diversity analyses, MaAsLin, and FAPROTAX).

**Supplementary Figure 1.**
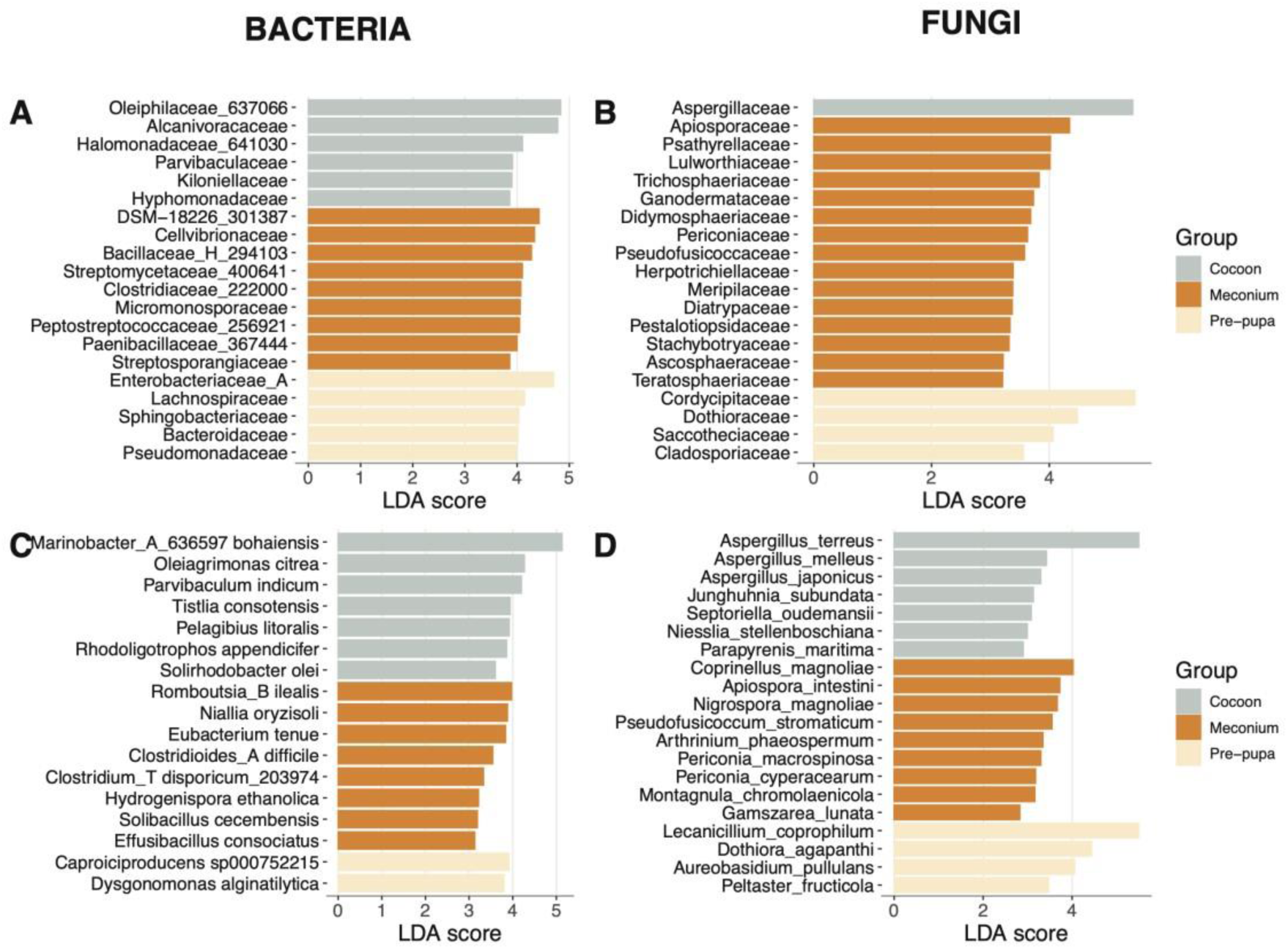
**Linear discriminant analysis (LDA) effect size (LEfSe)** at the family (A and B) and species (C and D) levels of bacteria (A,C) and fungi (B,D) showing the top 20 significant taxa for each *Centris aethyctera’s* brood cell sample type (alpha=0.05, LDA score threshold=4). Unknown families and species are not shown in the LEfSe analysis.

**Supplementary Figure 2.**
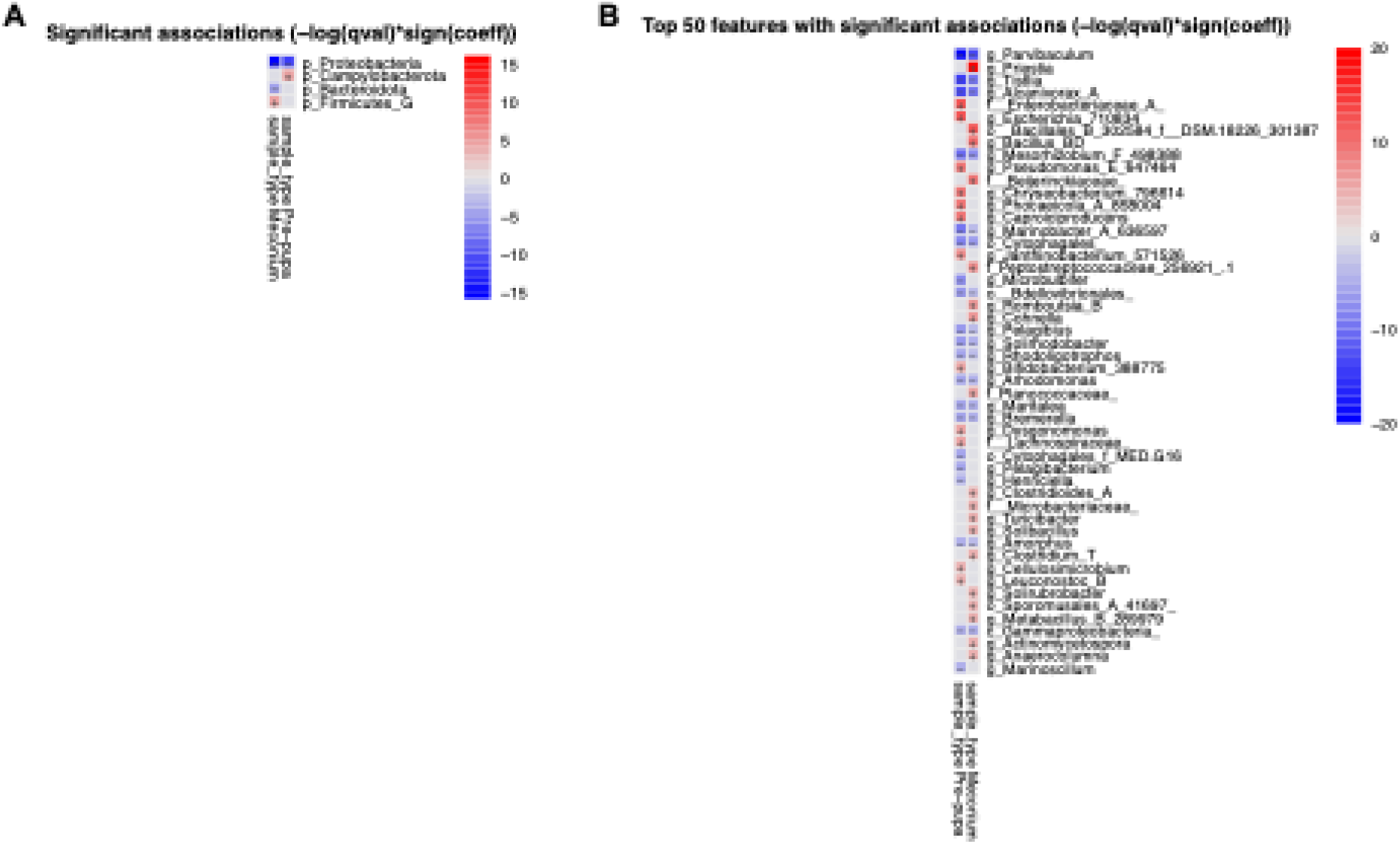
**Heatmaps from the bacterial Multivariate Analysis by Linear Models (MaAsLin) in each brood cell sample types (Cocoon, Meconium, and Pre-pupa) of *Centris aethyctera*** at the phylum (A) and genus level from which the top 50 features are shown (B). The reference point of sample type was set to Cocoon (so the taxa from other sample types being either more or less associated than Cocoon), as it has the most uniform bacterial composition. Red (+) indicates a positive association while blue (-) a negative one. P-value threshold is set at 0.01.

**Supplementary Figure 3.**
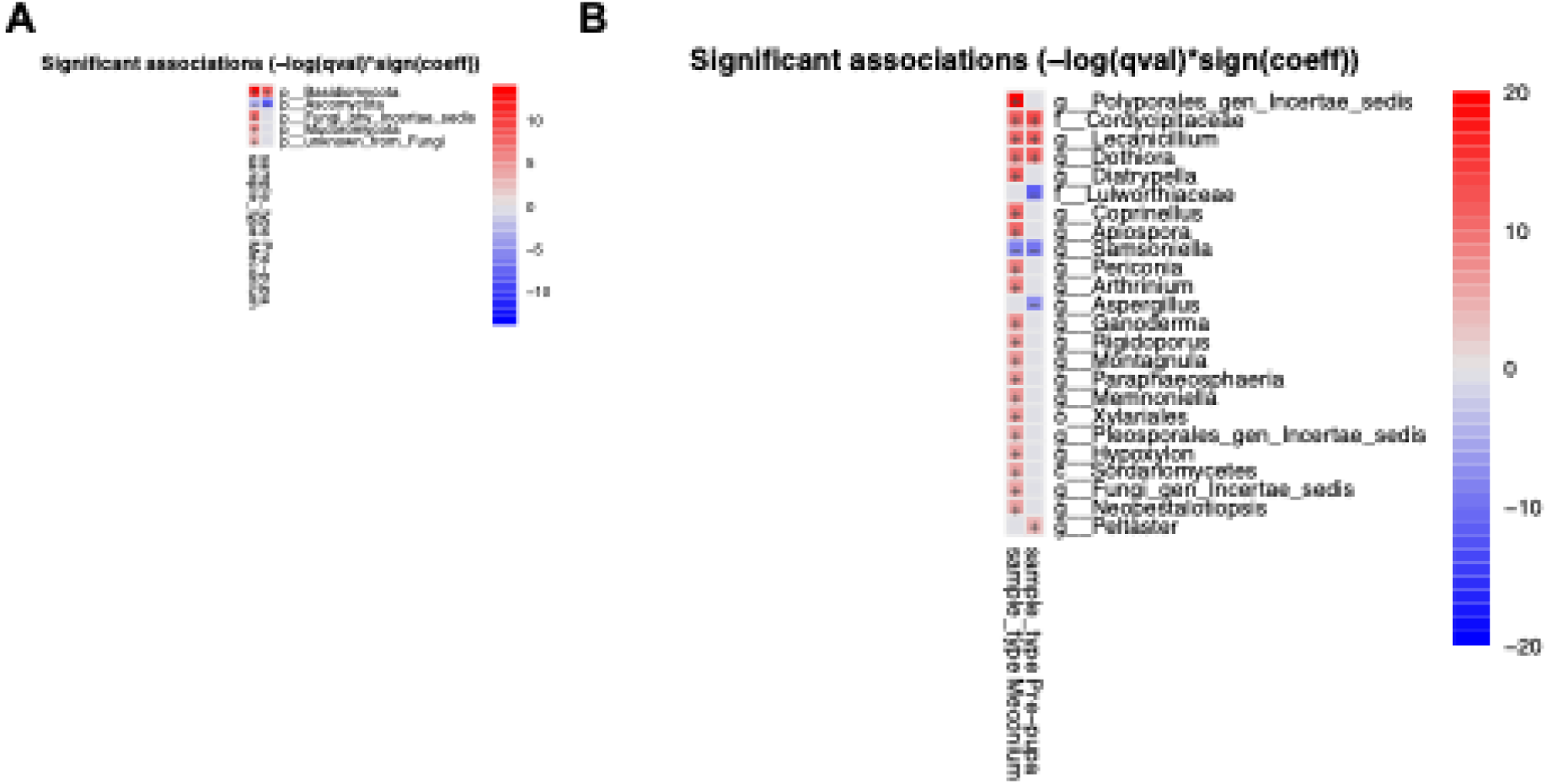
Results from the fungal Multivariate Analysis by Linear Models (MaAsLin) in each brood cell sample types (Cocoon, Meconium, and Pre-pupa) of *Centris aethyctera*: (A) Heatmaps at the phylum level with the reference point of sample type set to Cocoon (so the taxa from other sample types being either more or less associated than Cocoon), as it has the most uniform bacterial composition. Red (+) indicates a positive association while blue (-) a negative one; (B) Heatmaps at the genus level with the reference point of sample type set to Cocoon (so the taxa from other sample types being either more or less associated than Cocoon), as it has the most uniform bacterial composition. Red (+) indicates a positive association while blue (-) a negative one. P-value threshold is set at 0.01.

**Supplementary Figure 4.**
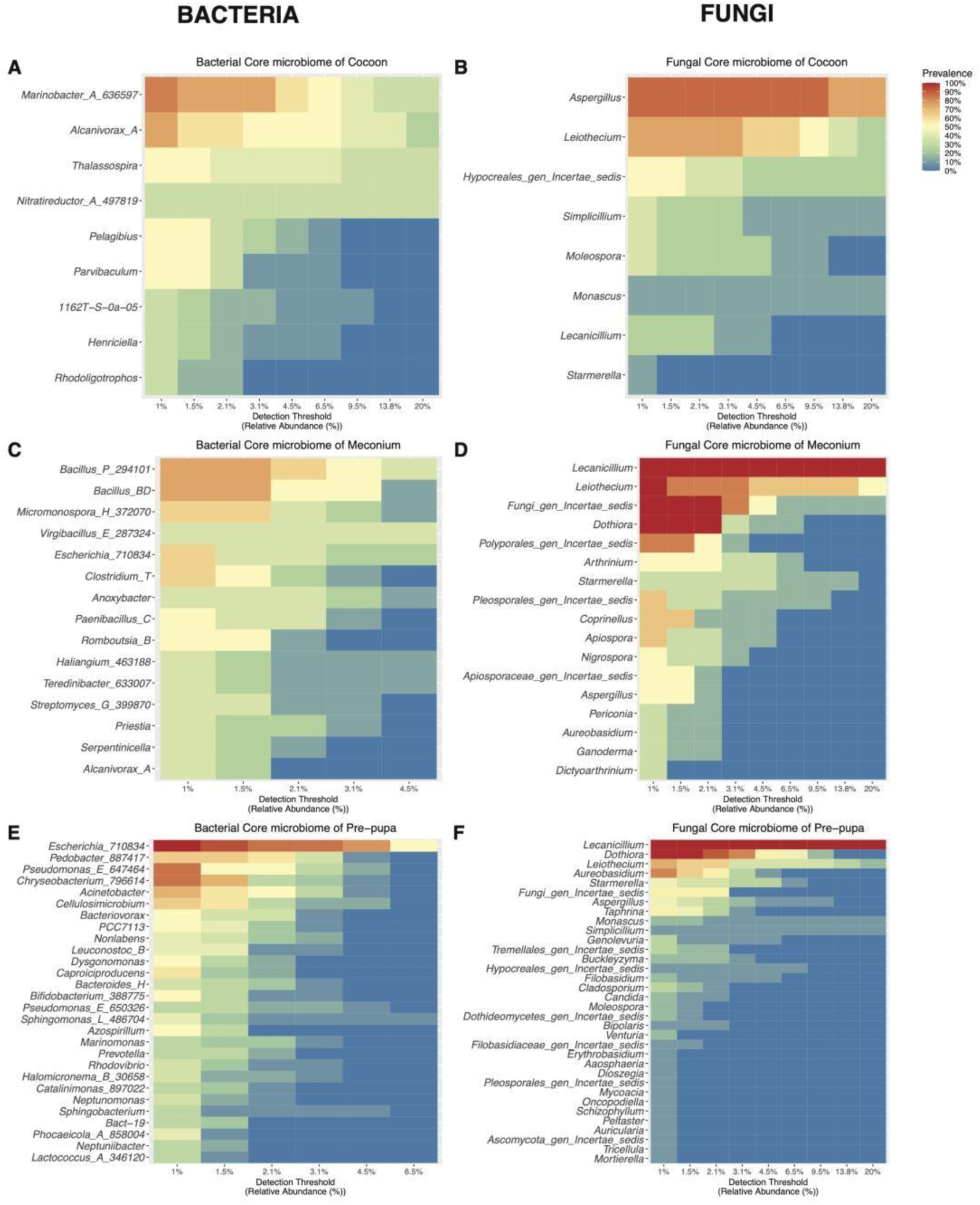
**Heatmaps of Core Microbiome of brood cells sample types of *Centris aethyctera* (Cocoon, Meconium, and Pre-pupa)** at the genus level (y axis). Colors indicates the prevalence of the taxa in the samples, from blue/0% (present in 0% of all the samples) to red/100% (present in 100% of the samples), with a certain detection threshold (x axis) indicating the relative prevalence in % of the taxa within the sample. Panels A, C, and E represent the heatmaps of bacterial core microbiome in either Cocoon, Meconium, or Prepupa, respectively, while Panels B, D, and F represent the heatmaps of fungal core microbiome in either Cocoon, Meconium, or Pre-pupa, respectively.

**Supplementary Figure 5.**
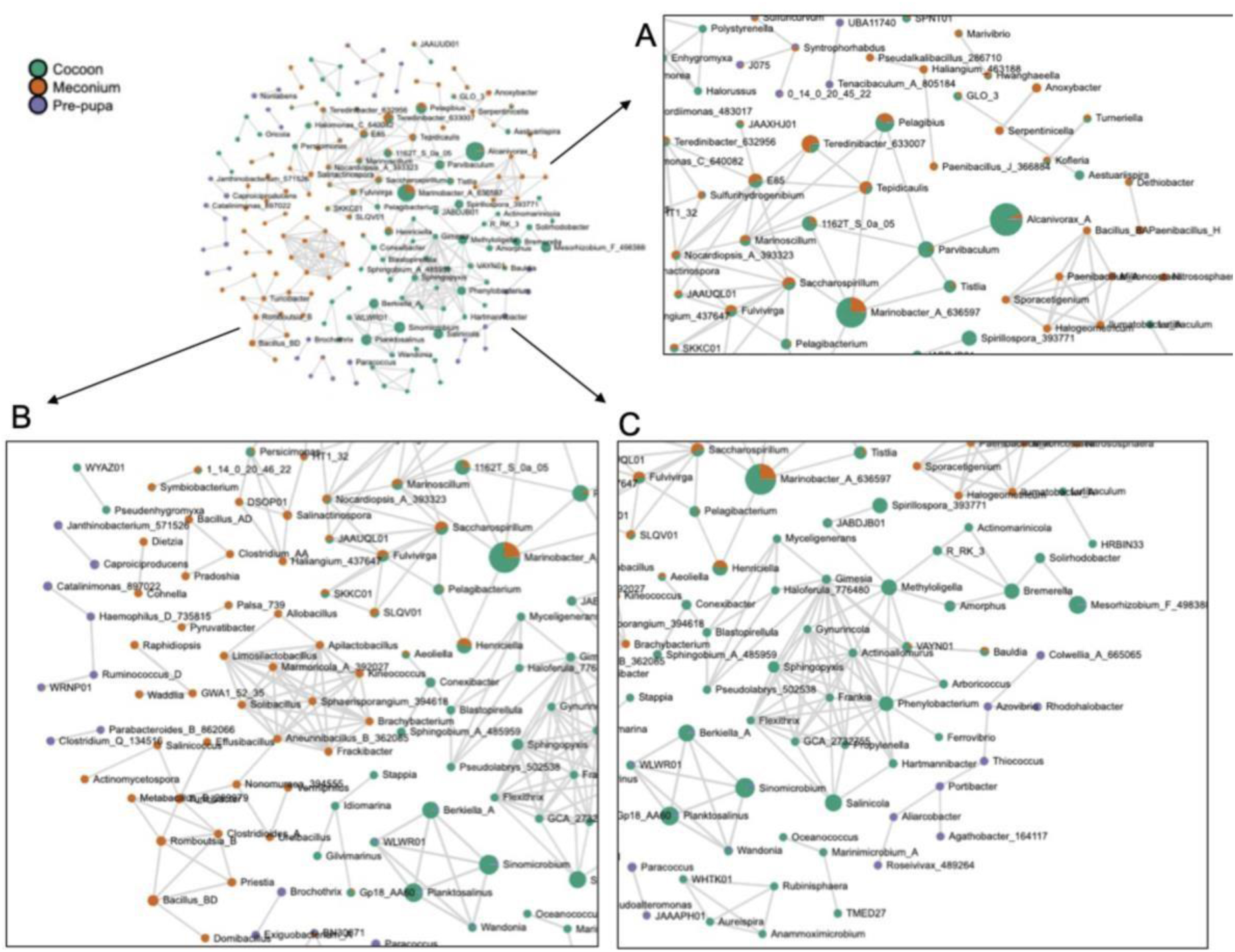
**Correlation Network Analysis of bacterial genus for each sample type of *Centris aethyctera’s* brood cells**, using SparCC method (parameters: SparCC, permutation=100, p-value threshold=0.01, correlation threshold=0.7, at the genus level). In green is shown the cocoon, in red the meconium, and in purple the pre-pupa. Each node represents the proportion of this bacteria in each sample type using a pie-chart. To compute this network in Microbiome Analyst 2.0, no count filter, no variance filter, no rarefaction, or no data transformation were used, but the data was scaled to the total sum scaling (TSS).

**Supplementary Figure 6.**
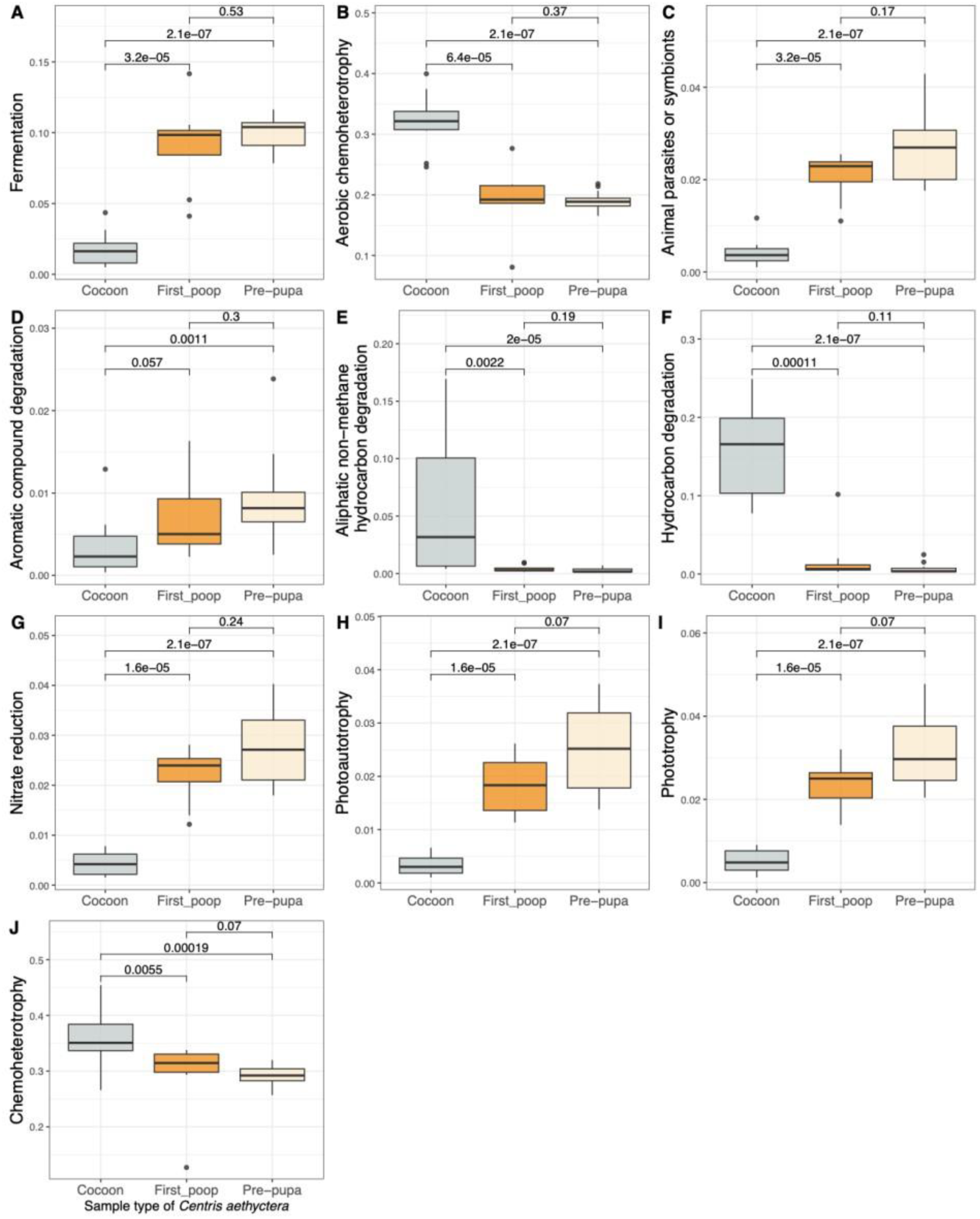
Boxplots showing the significant differences in bacterial functional activity. **(A-J) of each brood cell sample type** (Cocoon, Meconium, and Prepupa) of *Centris aethyctera* using Functional Activity of Prokaryote taxa (FAPROTAX) database. The functions shown are those who shown a standard deviation higher than 0.01 when comparing the relative abundance of functional active taxa between samples.

**Supplementary Figure 7.**
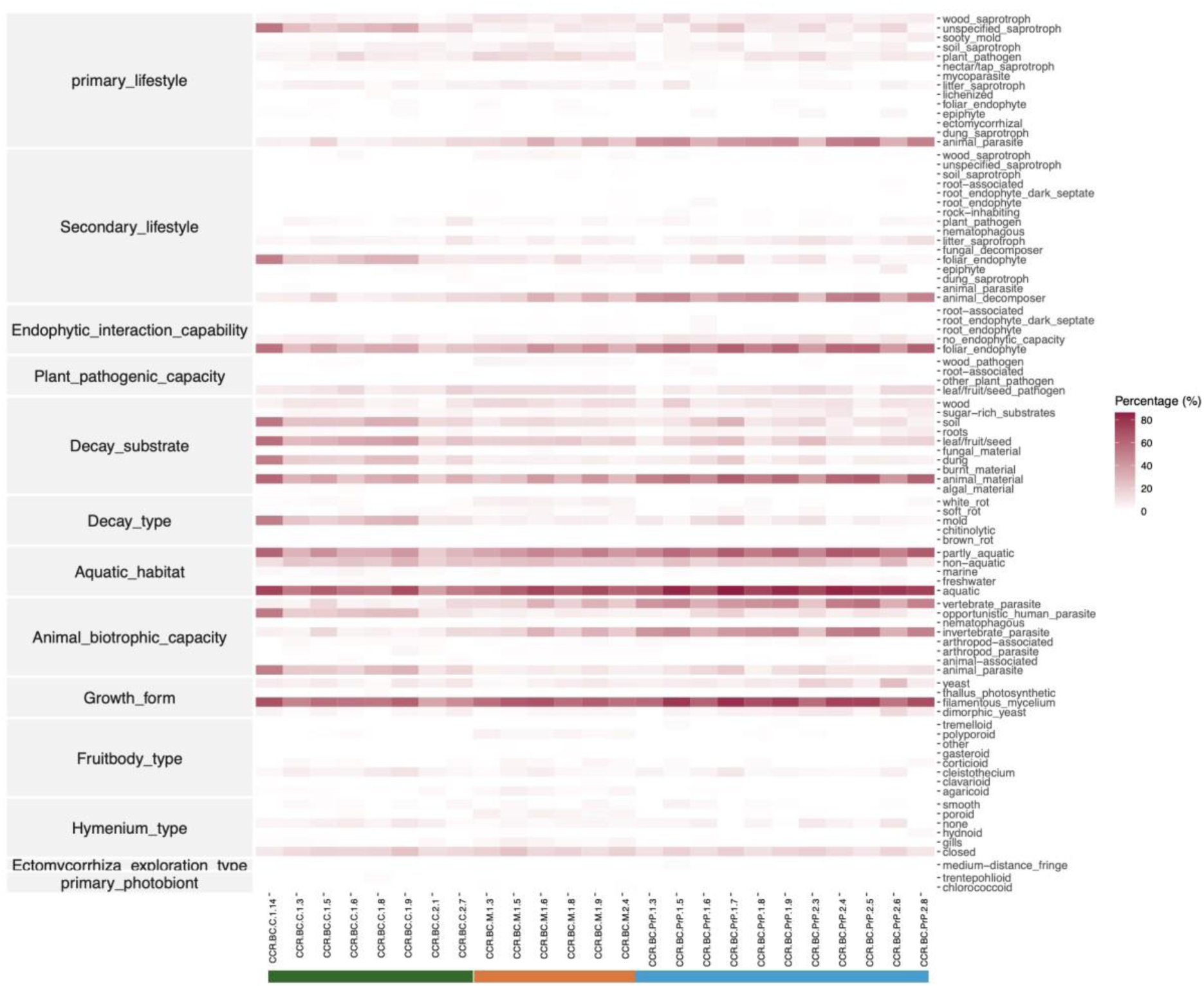
Heatmap showing the different fungal functions divided in categories. (left side of the heatmap): primary and secondary lifestyles, endophytic interaction capability, plant pathogenic capacity, decay substrate, decay type, aquatic habitat, animal biotrophic capacity, growth form, fruitbody type, hymenium type, Ectomycorrhiza exploration type, and primary photobiont). Details for each category are shown at the right side of the heatmap. Sample IDs are shown on the x axis, with the color bar representing Cocoon (grey), Meconium (orange), Pre-pupa (yellow). The abundance of each function per sample type (in %) is shown in a gradient of red.

**Supplementary Figure 8.**
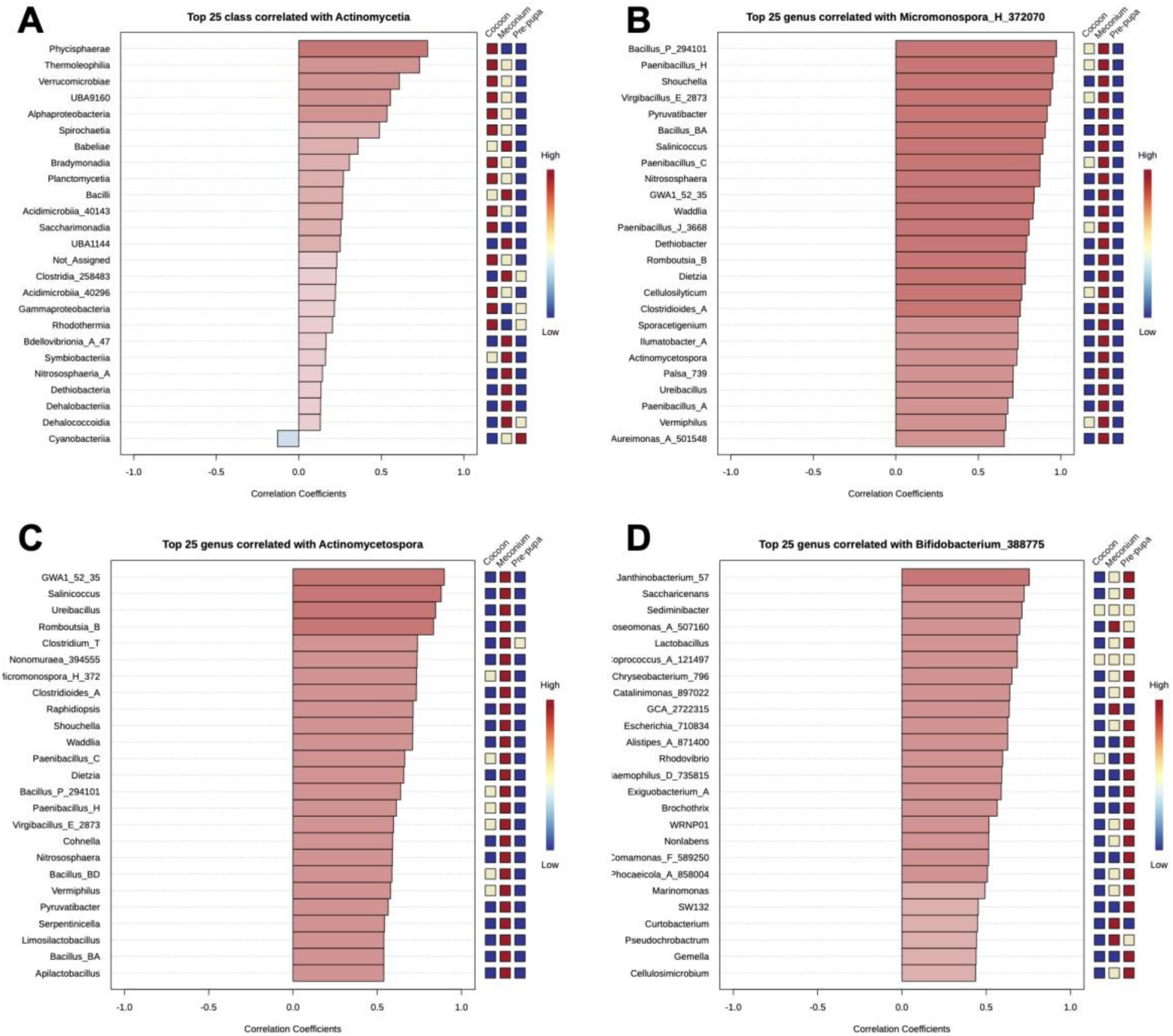
**Results of Pattern Search** from Microbiome Analyst 2.0, using Pearson r correlation index, showing the top 25 bacteria positively correlated (positive correlation coefficient shades of red) or negatively correlated (negative correlation coefficient shades of blue) with (A) Actinomycetia, and biomarkers of meconium samples, i.e., *Micromonospora, Actinomycetospora*, and biomarkers of pre-pupa samples, i.e., *Bifidobacterium*. The sample types of brood cells (Cocoon, Meconium, and Pre-pupa) from *Centris aethyctera* are shown on the right side of each graph with 3 levels of colors: ‘red’ represents taxa being highly correlated to this sample, ‘yellowish white’ represents taxa not being correlated either positively or negatively, and ‘blue’ represents the negative correlation to this sample. The scale of correlation ranging from high to low is shown on the very right side of each graph.

**Supplementary Figure 9.**
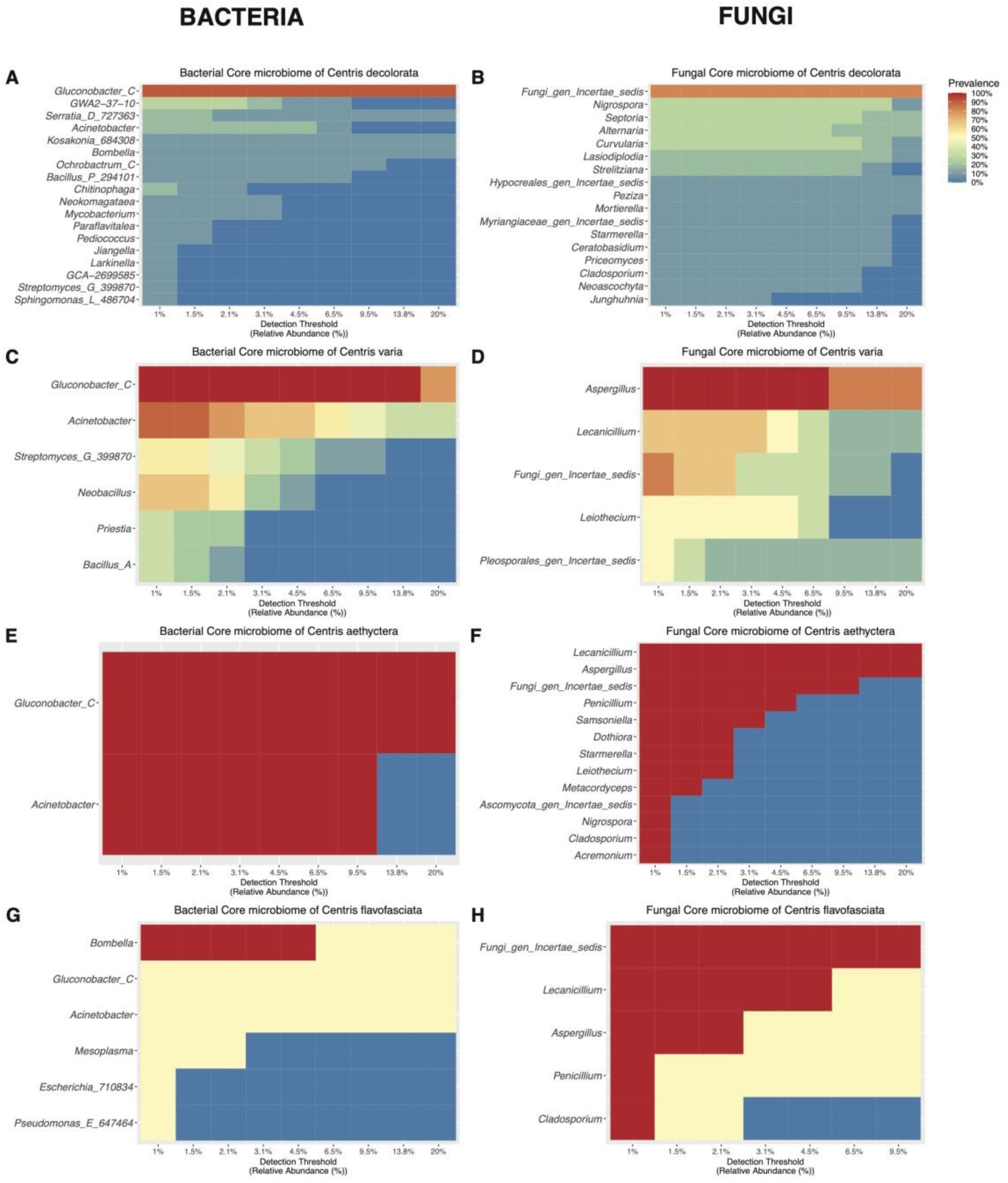
Heatmaps of Core Microbiome among female Centris guts per species, at the genus level (y axis). Colors indicates the prevalence of the taxa in the samples, from blue/0% (present in 0% of all the samples) to red/100% (present in 100% of the samples), with a certain detection threshold (x axis) indicating the relative prevalence in % of the taxa within the sample. Panels A and B respectively represents the heatmaps of bacterial and fungal core microbiome in *Centris decolorata* (Puerto Rico), Panels C and D respectively represents the heatmaps of bacterial and fungal core microbiome in *Centris varia* (Costa Rica), Panels E and F respectively represents the heatmaps of bacterial and fungal core microbiome in *Centris aethyctera* (Costa Rica), Panels G and H respectively represents the heatmaps of bacterial and fungal core microbiome in *Centris flavofasciata* (Costa Rica).

## References

[1] Vannette RL, Gauthier M-PL, Fukami T. Nectar bacteria, but not yeast, weaken a plant–pollinator mutualism. Proceedings of the Royal Society B: Biological Sciences 2013;280:20122601. 10.1098/rspb.2012.2601.

[2] Steffan SA, Dharampal PS, Danforth BN, Gaines-Day HR, Takizawa Y, Chikaraishi Y. Omnivory in Bees: Elevated Trophic Positions among All Major Bee Families. Am Nat 2019;194:414–21. 10.1086/704281.

[3] McFrederick QS, Mueller UG, James RR. Interactions between fungi and bacteria influence microbial community structure in the *Megachile rotundata* larval gut. Proceedings of the Royal Society B: Biological Sciences 2014;281:20132653. 10.1098/rspb.2013.2653.

[4] Kaltenpoth M, Engl T. Defensive microbial symbionts in Hymenoptera. Funct Ecol 2014;28:315–27. 10.1111/1365-2435.12089.

[5] Raymann K, Moran NA. The role of the gut microbiome in health and disease of adult honey bee workers. Curr Opin Insect Sci 2018;26:97–104. 10.1016/j.cois.2018.02.012.

[6] Voulgari-Kokota A, Grimmer G, Steffan-Dewenter I, Keller A. Bacterial community structure and succession in nests of two megachilid bee genera. FEMS Microbiol Ecol 2019;95. 10.1093/femsec/fiy218.

[7] Voulgari-Kokota A, Ankenbrand MJ, Grimmer G, Steffan-Dewenter I, Keller A. Linking pollen foraging of megachilid bees to their nest bacterial microbiota. Ecol Evol 2019;9:10788–800. 10.1002/ece3.5599.

[8] Rothman JA, Andrikopoulos C, Cox-Foster D, McFrederick QS. Floral and Foliar Source Affect the Bee Nest Microbial Community. Microb Ecol 2019;78:506–16. 10.1007/s00248-018-1300-3.

[9] Dharampal PS, Hetherington MC, Steffan SA. Microbes make the meal: oligolectic bees require microbes within their host pollen to thrive. Ecol Entomol 2020;45:1418–27. 10.1111/een.12926.

[10] McFrederick QS, Rehan SM. Characterization of pollen and bacterial community composition in brood provisions of a small carpenter bee. Mol Ecol 2016;25:2302–11. 10.1111/mec.13608.

[11] McFrederick QS, Thomas JM, Neff JL, Vuong HQ, Russell KA, Hale AR, et al. Flowers and Wild Megachilid Bees Share Microbes. Microb Ecol 2017;73:188–200. 10.1007/s00248-016-0838-1.

[12] Voulgari-Kokota A, McFrederick QS, Steffan-Dewenter I, Keller A. Drivers, Diversity, and Functions of the Solitary-Bee Microbiota. Trends Microbiol 2019;27:1034–44. 10.1016/j.tim.2019.07.011.

[13] Voulgari-Kokota A, Ankenbrand MJ, Grimmer G, Steffan-Dewenter I, Keller A. Linking pollen foraging of megachilid bees to their nest bacterial microbiota. Ecol Evol 2019;9:10788–800. 10.1002/ece3.5599.

[14] Voulgari-Kokota A, Steffan-Dewenter I, Keller A. Susceptibility of Red Mason Bee Larvae to Bacterial Threats Due to Microbiome Exchange with Imported Pollen Provisions. Insects 2020;11:373. 10.3390/insects11060373.

[15] Danforth BN, Minckley RL, Neff JL, Fawcett F. The Solitary Bees. Princeton University Press; 2019. 10.1515/9780691189321.

[16] Flórez L V., Scherlach K, Gaube P, Ross C, Sitte E, Hermes C, et al. Antibiotic-producing symbionts dynamically transition between plant pathogenicity and insect-defensive mutualism. Nat Commun 2017;8:15172. 10.1038/ncomms15172.

[17] Flórez L V., Biedermann PHW, Engl T, Kaltenpoth M. Defensive symbioses of animals with prokaryotic and eukaryotic microorganisms. Nat Prod Rep 2015;32:904–36. 10.1039/C5NP00010F.

[18] Chevrette MG, Carlson CM, Ortega HE, Thomas C, Ananiev GE, Barns KJ, et al. The antimicrobial potential of *Streptomyces* from insect microbiomes. Nat Commun 2019;10:516. 10.1038/s41467-019-08438-0.

[19] Fernández-Marín H, Zimmerman JK, Nash DR, Boomsma JJ, Wcislo WT. Reduced biological control and enhanced chemical pest management in the evolution of fungus farming in ants. Proceedings of the Royal Society B: Biological Sciences 2009;276:2263–9. 10.1098/rspb.2009.0184.

[20] Kaltenpoth M, Göttler W, Herzner G, Strohm E. Symbiotic Bacteria Protect Wasp Larvae from Fungal Infestation. Current Biology 2005;15:475–9. 10.1016/j.cub.2004.12.084.

[21] Hammer TJ, Kueneman J, Argueta-Guzmán M, McFrederick QS, Grant, Lady, Wcislo W, et al. Bee breweries: The unusually fermentative, lactobacilli-dominated brood cell microbiomes of cellophane bees. Front Microbiol 2023;14. 10.3389/fmicb.2023.1114849.

[22] Flórez L V., Biedermann PHW, Engl T, Kaltenpoth M. Defensive symbioses of animals with prokaryotic and eukaryotic microorganisms. Nat Prod Rep 2015;32:904–36. 10.1039/C5NP00010F.

[23] Poulsen M, Cafaro M, Boomsma JJ, Currie CR. Specificity of the mutualistic association between actinomycete bacteria and two sympatric species of *Acromyrmex* leaf-cutting ants. Mol Ecol 2005;14:3597–604. 10.1111/j.1365-294X.2005.02695.x.

[24] Mueller UG, Dash D, Rabeling C, Rodrigues A. Coevolution between attine ants and actinomycete bacteria: a reevaluation. Evolution (N Y) 2008;62:2894–912. 10.1111/j.1558-5646.2008.00501.x.

[25] Cafaro MJ, Poulsen M, Little AEF, Price SL, Gerardo NM, Wong B, et al. Specificity in the symbiotic association between fungus-growing ants and protective *Pseudonocardia* bacteria. Proceedings of the Royal Society B: Biological Sciences 2011;278:1814–22. 10.1098/rspb.2010.2118.

[26] Madden AA, Grassetti A, Soriano J-AN, Starks PT. *Actinomycetes* with Antimicrobial Activity Isolated from Paper Wasp (Hymenoptera: Vespidae: Polistinae) Nests. Environ Entomol 2013;42:703–10. 10.1603/EN12159.

[27] Kaltenpoth M, Yildirim E, Gürbüz MF, Herzner G, Strohm E. Refining the Roots of the Beewolf-Streptomyces Symbiosis: Antennal Symbionts in the Rare Genus Philanthinus (Hymenoptera, Crabronidae). Appl Environ Microbiol 2012;78:822–7. 10.1128/AEM.06809-11.

[28] Kaltenpoth M, Goettler W, Dale C, Stubblefield JW, Herzner G, Roeser-Mueller K, et al. ‘Candidatus *Streptomyces philanthi*’, an endosymbiotic streptomycete in the antennae of Philanthus digger wasps. Int J Syst Evol Microbiol 2006;56:1403–11. 10.1099/ijs.0.64117-0.

[29] Kaltenpoth M, Roeser-Mueller K, Koehler S, Peterson A, Nechitaylo TY, Stubblefield JW, et al. Partner choice and fidelity stabilize coevolution in a Cretaceous-age defensive symbiosis. Proceedings of the National Academy of Sciences 2014;111:6359–64. 10.1073/pnas.1400457111.

[30] Engl T, Kroiss J, Kai M, Nechitaylo TY, Svatoš A, Kaltenpoth M. Evolutionary stability of antibiotic protection in a defensive symbiosis. Proceedings of the National Academy of Sciences 2018;115. 10.1073/pnas.1719797115.

[31] Cambronero-Heinrichs JC, Matarrita-Carranza B, Murillo-Cruz C, Araya-Valverde E, Chavarría M, Pinto-Tomás AA. Phylogenetic analyses of antibiotic-producing *Streptomyces* sp. isolates obtained from the stingless-bee *Tetragonisca angustula* (Apidae: Meliponini). Microbiology (N Y) 2019;165:292–301. 10.1099/mic.0.000754.

[32] Promnuan Y, Kudo T, Chantawannakul P. *Actinomycetes* isolated from beehives in Thailand. World J Microbiol Biotechnol 2009;25:1685–9. 10.1007/s11274-009-0051-1.

[33] Grubbs KJ, May DS, Sardina JA, Dermenjian RK, Wyche TP, Pinto-Tomás AA, et al. Pollen Streptomyces Produce Antibiotic That Inhibits the Honey Bee Pathogen *Paenibacillus larvae*. Front Microbiol 2021;12. 10.3389/fmicb.2021.632637.

[34] Kardas E, González-Rosario AM, Giray T, Ackerman JD, Godoy-Vitorino F. Gut microbiota variation of a tropical oil-collecting bee species far exceeds that of the honeybee. Front Microbiol 2023;14. 10.3389/fmicb.2023.1122489.

[35] Hanson PE, Fernández-Otárola M, Lobo Segura J, Frankie GW, Coville R, Aguilar Monge I, et al. Bees of Costa Rica. Ithaca and London: Cornell University Press; 2023.

[36] Roubik DW, editor. Ecology and Natural History of Tropical Bees. Cambridge University Press; 1989. 10.1017/CBO9780511574641.

[37] Keller A, Grimmer G, Steffan-Dewenter I. Diverse Microbiota Identified in Whole Intact Nest Chambers of the Red Mason Bee *Osmia bicornis* (Linnaeus 1758). PLoS One 2013;8:e78296. 10.1371/journal.pone.0078296.

[38] Lobo JA, Fernández Otárola M, Chavarría MM, Agraz Hernández CM. Nesting biology of *Centris aethyctera* (Centridini, Apidae) in an estuarine environment. Apidologie 2023;54:1–14. 10.1007/s13592-023-01044-6.

[39] Kardas E, González-Rosario AM, Giray T, Ackerman JD, Godoy-Vitorino F. Gut microbiota variation of a tropical oil-collecting bee species far exceeds that of the honeybee. Front Microbiol 2023;14. 10.3389/fmicb.2023.1122489.

[40] McDonald D, Jiang Y, Balaban M, Cantrell K, Zhu Q, Gonzalez A, et al. Greengenes2 unifies microbial data in a single reference tree. Nat Biotechnol 2023. 10.1038/s41587-02301845-1.

[41] Bolyen E, Rideout JR, Dillon MR, Bokulich NA, Abnet CC, Al-Ghalith GA, et al. Reproducible, interactive, scalable and extensible microbiome data science using QIIME 2. Nat Biotechnol 2019;37:852–7. 10.1038/s41587-019-0209-9.

[42] Quast C, Pruesse E, Yilmaz P, Gerken J, Schweer T, Yarza P, et al. The SILVA ribosomal RNA gene database project: improved data processing and web-based tools. Nucleic Acids Res 2012;41:D590–6. 10.1093/nar/gks1219.

[43] Wickham H, François R, Henry L, Müller K, Vaughan D. dplyr: A Grammar of Data Manipulation. 2023.

[44] Abarenkov K, Nilsson RH, Larsson KH, Taylor AFS, May TW, Frøslev TG, et al. The UNITE database for molecular identification and taxonomic communication of fungi nd other eukaryotes: sequences, taxa and classifications reconsider ed. Nucleic Acids Res 2024;52:D791–7. 10.1093/nar/gkad1039.

[45] Rodríguez-Barreras R, Tosado-Rodríguez EL, Godoy-Vitorino F. Trophic niches reflect compositional differences in microbiota among Caribbean sea urchins. PeerJ 2021;9:e12084. 10.7717/peerj.12084.

[46] Ruiz Barrionuevo JM, Vilanova-Cuevas B, Alvarez A, Martín E, Malizia A, Galindo-Cardona A, et al. The Bacterial and Fungal Gut Microbiota of the Greater Wax Moth, *Galleria mellonella* L. Consuming Polyethylene and Polystyrene. Front Microbiol 2022;13. 10.3389/fmicb.2022.918861.

[47] Kardas E, González-Rosario AM, Giray T, Ackerman JD, Godoy-Vitorino F. Gut microbiota variation of a tropical oil-collecting bee species far exceeds that of the honeybee. Front Microbiol 2023;14. 10.3389/fmicb.2023.1122489.

[48] Faith DP. Conservation evaluation and phylogenetic diversity. Biol Conserv 1992;61:1–10. 10.1016/0006-3207(92)91201-3.

[49] Shannon CE. A Mathematical Theory of Communication. Bell System Technical Journal 1948;27:379–423. 10.1002/j.1538-7305.1948.tb01338.x.

[50] Liu C, Cui Y, Li X, Yao M. microeco: an R package for data mining in microbial community ecology. FEMS Microbiol Ecol 2021;97. 10.1093/femsec/fiaa255.

[51] Kruskal WH, Wallis WA. Use of Ranks in One-Criterion Variance Analysis. J Am Stat Assoc 1952;47:583. 10.2307/2280779.

[52] Anderson MJ. A new method for non-parametric multivariate analysis of variance. Austral Ecol 2001;26:32–46. 10.1111/j.1442-9993.2001.01070.pp.x.

[53] Clarke KR. Non-parametric multivariate analyses of changes in community structure. Australian Journal of Ecology 1993;18:117–43. 10.1111/j.1442-9993.1993.tb00438.x.

[54] McArdle BH, Anderson MJ. Fitting Multivariate Models to Community Data: A Comment on Distance-Based Redundancy Analysis. Ecology 2001;82. 10.1890/00129658(2001)082[0290:FMMTCD]2.0.CO;2.

[55] McMurdie PJ, Holmes S. phyloseq: An R Package for Reproducible Interactive Analysis and Graphics of Microbiome Census Data. PLoS One 2013;8:e61217. 10.1371/journal.pone.0061217.

[56] Oksanen J, Simpson G, Blanchet F, Kindt R, Legendre P, Minchin P, et al. vegan: Community Ecology Package. R package version 2.6–5 2024.

[57] Wickham H. ggplot2: Elegant Graphics for Data Analysis. 2016.

[58] Mallick H, Rahnavard A, McIver LJ, Ma S, Zhang Y, Nguyen LH, et al. Multivariable association discovery in population-scale meta-omics studies. PLoS Comput Biol 2021;17:e1009442. 10.1371/journal.pcbi.1009442.

[59] Segata N, Izard J, Waldron L, Gevers D, Miropolsky L, Garrett WS, et al. Metagenomic biomarker discovery and explanation. Genome Biol 2011;12:R60. 10.1186/gb-2011-126-r60.

[60] Biomarkers Definitions Working Group. Biomarkers and surrogate endpoints: Preferred definitions and conceptual framework. Clin Pharmacol Ther 2001;69:89–95. 10.1067/mcp.2001.113989.

[61] Strimbu K, Tavel JA. What are biomarkers? Curr Opin HIV AIDS 2010;5:463–6. 10.1097/COH.0b013e32833ed177.

[62] Huang H-C, Jupiter D, VanBuren V. Classification of Genes and Putative Biomarker Identification Using Distribution Metrics on Expression Profiles. PLoS One 2010;5:e9056. 10.1371/journal.pone.0009056.

[63] Friedman J, Alm EJ. Inferring Correlation Networks from Genomic Survey Data. PLoS Comput Biol 2012;8:e1002687. 10.1371/journal.pcbi.1002687.

[64] Lu Y, Zhou G, Ewald J, Pang Z, Shiri T, Xia J. MicrobiomeAnalyst 2.0: comprehensive statistical, functional and integrative analysis of microbiome data. Nucleic Acids Res 2023;51:W310–8. 10.1093/nar/gkad407.

[65] Louca S, Parfrey LW, Doebeli M. Decoupling function and taxonomy in the global ocean microbiome. Science (1979) 2016;353:1272–7. 10.1126/science.aaf4507.

[66] Põlme S, Abarenkov K, Henrik Nilsson R, Lindahl BD, Clemmensen KE, Kauserud H, et al. FungalTraits: a user-friendly traits database of fungi and fungus-like stramenopiles. Fungal Divers 2020;105:1–16. 10.1007/s13225-020-00466-2.

[67] McMurdie PJ, Holmes S. Waste Not, Want Not: Why Rarefying Microbiome Data Is Inadmissible. PLoS Comput Biol 2014;10:e1003531. 10.1371/journal.pcbi.1003531.

[68] Gomez-Escribano J, Alt S, Bibb M. Next Generation Sequencing of Actinobacteria for the Discovery of Novel Natural Products. Mar Drugs 2016;14:78. 10.3390/md14040078.

[69] Liu C, Shao Z. *Alcanivorax dieselolei* sp. nov., a novel alkane-degrading bacterium isolated from sea water and deep-sea sediment. Int J Syst Evol Microbiol 2005;55:1181–6. 10.1099/ijs.0.63443-0.

[70] Yang S-H, Seo H-S, Seong CN, Kwon KK. *Oleiagrimonas citrea* sp. nov., a marine bacterium isolated from tidal flat sediment and emended description of the genus *Oleiagrimonas* Fang et al. 2015 and *Oleiagrimonas* soli. Int J Syst Evol Microbiol 2017;67:1672–5. 10.1099/ijsem.0.001846.

[71] Lobo JA, Fernández Otárola M, Chavarría MM, Agraz Hernández CM. Nesting biology of *Centris aethyctera* (Centridini, Apidae) in an estuarine environment. Apidologie 2023;54. 10.1007/s13592-023-01044-6.

[72] Middelburg JJ. Carbon Processing at the Seafloor, 2019, p. 57–75. 10.1007/9783-030-10822-9_4.

[73] Ettwig KF, Butler MK, Le Paslier D, Pelletier E, Mangenot S, Kuypers MMM, et al. Nitritedriven anaerobic methane oxidation by oxygenic bacteria. Nature 2010;464:543–8. 10.1038/nature08883.

[74] Kumar N, Srivastava P, Vishwakarma K, Kumar R, Kuppala H, Maheshwari SK, et al. The Rhizobium–Plant Symbiosis: State of the Art. Plant Microbe Symbiosis, Cham: Springer International Publishing; 2020, p. 1–20. 10.1007/978-3-030-36248-5_1.

[75] Chu C, Liu B, Lian Z, Zheng H, Chen C, Yue Z, et al. *Solirhodobacter olei* gen. nov., sp. nov., a nonphotosynthetic bacterium isolated from oil-contaminated soil. Int J Syst Evol Microbiol 2020;70:582–8. 10.1099/ijsem.0.003795.

[76] Choi DH, Hwang CY, Cho BC. *Pelagibius litoralis* gen. nov., sp. nov., a marine bacterium in the family Rhodospirillaceae isolated from coastal seawater. Int J Syst Evol Microbiol 2009;59:818–23. 10.1099/ijs.0.002774-0.

[77] Gardner JG. Polysaccharide degradation systems of the saprophytic bacterium *Cellvibrio japonicus*. World J Microbiol Biotechnol 2016;32:121. 10.1007/s11274-016-2068-6.

[78] Qiao N, Shao Z. Isolation and characterization of a novel biosurfactant produced by hydrocarbon-degrading bacterium *Alcanivorax dieselolei* B-5. J Appl Microbiol 2010;108:1207–16. 10.1111/j.1365-2672.2009.04513.x.

[79] Friedlander A, Nir S, Reches M, Shemesh M. Preventing Biofilm Formation by DairyAssociated Bacteria Using Peptide-Coated Surfaces. Front Microbiol 2019;10. 10.3389/fmicb.2019.01405.

[80] Shatalin K, Shatalina E, Mironov A, Nudler E. H 2 S: A Universal Defense Against Antibiotics in Bacteria. Science (1979) 2011;334:986–90. 10.1126/science.1209855.

[81] Hirsch AM, Valdés M. *Micromonospora*: An important microbe for biomedicine and potentially for biocontrol and biofuels. Soil Biol Biochem 2010;42:536–42. 10.1016/j.soilbio.2009.11.023.

[82] Wagman GH, Gannon RD, Weinstein MJ. Production of Vitamin B 12 by *Micromonospora*. Appl Microbiol 1969;17:648–9. 10.1128/am.17.4.648-649.1969.

[83] Ismet A, Vikineswary S, Paramaswari S, Wong WH, Ward A, Seki T, et al. Production and Chemical Characterization of Antifungal Metabolites From *Micromonospora* sp. M39 Isolated From Mangrove Rhizosphere Soil. World J Microbiol Biotechnol 2004;20:523–8. 10.1023/B:WIBI.0000040399.60343.4c.

[84] Rodríguez-Hernández D, Melo WGP, Menegatti C, Lourenzon VB, do Nascimento FS, Pupo MT. Actinobacteria associated with stingless bees biosynthesize bioactive polyketides against bacterial pathogens. New Journal of Chemistry 2019;43:10109–17. 10.1039/C9NJ01619H.

[85] Han T-Y, Tong X-M, Wang Y-W, Wang H-M, Chen X-R, Kong D-L, et al. *Paenibacillus populi* sp. nov., a novel bacterium isolated from the rhizosphere of *Populus alba*. Antonie Van Leeuwenhoek 2015;108:659–66. 10.1007/s10482-015-0521-4.

[86] Keller A, Brandel A, Becker MC, Balles R, Abdelmohsen UR, Ankenbrand MJ, et al. Wild bees and their nests host *Paenibacillus* bacteria with functional potential of avail. Microbiome 2018;6:229. 10.1186/s40168-018-0614-1.

[87] Keller A, Grimmer G, Steffan-Dewenter I. Diverse Microbiota Identified in Whole Intact Nest Chambers of the Red Mason Bee *Osmia bicornis* (Linnaeus 1758). PLoS One 2013;8:e78296. 10.1371/journal.pone.0078296.

[88] Lozo J, Berić T, Terzić-Vidojević A, Stanković S, Fira D, Stanisavljević L. Microbiota associated with pollen, bee bread, larvae and adults of solitary bee *Osmia cornuta* (Hymenoptera: Megachilidae). Bull Entomol Res 2015;105:470–6. 10.1017/S0007485315000292.

[89] Girardin H, Albagnac C, Dargaignaratz C, Nguyen-The C, Carlin F. Antimicrobial Activity of Foodborne *Paenibacillus* and *Bacillus* spp. against *Clostridium botulinum*. J Food Prot 2002;65:806– 13. 10.4315/0362-028X-65.5.806.

[90] Weid I, Alviano DS, Santos ALS, Soares RMA, Alviano CS, Seldin L. Antimicrobial activity of *Paenibacillus peoriae* strain NRRL BD-62 against a broad spectrum of phytopathogenic bacteria and fungi. J Appl Microbiol 2003;95:1143–51. 10.1046/j.1365-2672.2003.02097.x.

[91] Grady EN, MacDonald J, Liu L, Richman A, Yuan Z-C. Current knowledge and perspectives of *Paenibacillus*: a review. Microb Cell Fact 2016;15:203. 10.1186/s12934-016-0603-7.

[92] Nakano MM, Zuber P. Molecular Biology of Antibiotic Production in Bacillus. Crit Rev Biotechnol 1990;10:223–40. 10.3109/07388559009038209.

[93] Wahyudi AT, Prasojo BJ, Mubarik NR. Diversity of Antifungal Compounds-Producing *Bacillus* spp. Isolated from Rhizosphere of Soybean Plant Based on ARDRA and 16S rRNA. Hayati 2010;17:145–50. 10.4308/hjb.17.3.145.

[94] Guo B, Tang J, Ding G, Mashilingi SK, Huang J, An J. Gut microbiota is a potential factor in shaping phenotypic variation in larvae and adults of female bumble bees. Front Microbiol 2023;14. 10.3389/fmicb.2023.1117077.

[95] Dong Z-X, Li H-Y, Chen Y-F, Wang F, Deng X-Y, Lin L-B, et al. Colonization of the gut microbiota of honey bee (*Apis mellifera*) workers at different developmental stages. Microbiol Res 2020;231:126370. 10.1016/j.micres.2019.126370.

[96] Dai P, Wang M, Geng L, Yan Z, Yang Y, Guo L, et al. The effect of Bt Cry9Ee toxin on honey bee brood and adults reared in vitro, *Apis mellifera* (Hymenoptera: Apidae). Ecotoxicol Environ Saf 2019;181:381–7. 10.1016/j.ecoenv.2019.06.031.

[97] Cohen H, McFrederick QS, Philpott SM. Environment Shapes the Microbiome of the Blue Orchard Bee, *Osmia lignaria*. Microb Ecol 2020;80:897–907. 10.1007/s00248-02001549-y.

[98] Yong H-S, Song S-L, Chua K-O, Lim P-E. High Diversity of Bacterial Communities in Developmental Stages of *Bactrocera carambolae* (Insecta: Tephritidae) Revealed by Illumina MiSeq Sequencing of 16S rRNA Gene. Curr Microbiol 2017;74:1076–82. 10.1007/s00284-0171287-x.

[99] Vandamme P, Bernardet J-F, Segers P, Kersters K, Holmes B. NOTES: New Perspectives in the Classification of the Flavobacteria: Description of *Chryseobacterium* gen. nov., *Bergeyella* gen. nov., and *Empedobacter* nom. rev. Int J Syst Bacteriol 1994;44:827–31. 10.1099/00207713-44-4-827.

[100] Page AP, Roberts M, Félix M-A, Pickard D, Page A, Weir W. The golden death bacillus *Chryseobacterium nematophagum* is a novel matrix digesting pathogen of nematodes. BMC Biol 2019;17:10. 10.1186/s12915-019-0632-x.

[101] Mansfield MJ, Doxey AC. Genomic insights into the evolution and ecology of botulinum neurotoxins. Pathog Dis 2018;76. 10.1093/femspd/fty040.

[102] Aber RC, Wennersten C, Moellering RC. Antimicrobial Susceptibility of *Flavobacteria*. Antimicrob Agents Chemother 1978;14:483–7. 10.1128/AAC.14.3.483.

[103] Lim HJ, Shin HS. Antimicrobial and Immunomodulatory Effects of *Bifidobacterium* Strains: A Review. J Microbiol Biotechnol 2020;30:1793–800. 10.4014/jmb.2007.07046.

[104] Abdelhamid AG, Esaam A, Hazaa MM. Cell free preparations of probiotics exerted antibacterial and antibiofilm activities against multidrug resistant *E. coli*. Saudi Pharmaceutical Journal 2018;26:603–7. 10.1016/j.jsps.2018.03.004.

[105] Yang J, Yang H. Effect of Bifidobacterium breve in Combination With Different Antibiotics on *Clostridium difficile*. Front Microbiol 2018;9. 10.3389/fmicb.2018.02953.

[106] Dunlap P V., Urbanczyk H. Luminous Bacteria. The Prokaryotes, Berlin, Heidelberg: Springer Berlin Heidelberg; 2013, p. 495–528. 10.1007/978-3-642-30141-4_75.

[107] Menezes C, Vollet-Neto A, Marsaioli AJ, Zampieri D, Fontoura IC, Luchessi AD, et al. A Brazilian Social Bee Must Cultivate Fungus to Survive. Current Biology 2015;25:2851–5. 10.1016/j.cub.2015.09.028.

[108] Paludo CR, Menezes C, Silva-Junior EA, Vollet-Neto A, Andrade-Dominguez A, Pishchany G, et al. Stingless Bee Larvae Require Fungal Steroid to Pupate. Sci Rep 2018;8:1122. 10.1038/s41598-018-19583-9.

[109] Rutkowski D, Weston M, Vannette RL. Bees just wanna have fungi: a review of bee associations with nonpathogenic fungi. FEMS Microbiol Ecol 2023;99. 10.1093/femsec/fiad077.

[110] Atallah O, Yassin S. *Aspergillus* spp. eliminate *Sclerotinia sclerotiorum* by imbalancing the ambient oxalic acid concentration and parasitizing its sclerotia. Environ Microbiol 2020;22:5265–79. 10.1111/1462-2920.15213.

[111] Xu S, Wang D, Wei Y, Cui Q, Li W. *Marinobacter bohaiensis* sp. nov., a moderate halophile isolated from benthic sediment of the Bohai Sea. Int J Syst Evol Microbiol 2018;68:3534–9. 10.1099/ijsem.0.003025.

[112] Yuan B, Grau MF, Murata RM, Torok T, Venkateswaran K, Stajich JE, et al. Identification of the Neoaspergillic Acid Biosynthesis Gene Cluster by Establishing an In Vitro CRISPRRibonucleoprotein Genetic System in *Aspergillus melleus*. ACS Omega 2023;8:16713–21. 10.1021/acsomega.2c08104.

[113] Zhu F, Chen G, Chen X, Huang M, Wan X. Aspergicin, a new antibacterial alkaloid produced by mixed fermentation of two marine-derived mangrove epiphytic fungi. Chem Nat Compd 2011;47:767–9. 10.1007/s10600-011-0053-8.

[114] de Silva N. Morphomolecular taxonomic studies reveal a high number of endophytic fungi from *Magnolia candolli* and *M*. *garrettii* in China and Thailand. Mycosphere 2021;12:163–237. 10.5943/mycosphere/12/1/3.

[115] Baldrian P, Valášková V. Degradation of cellulose by basidiomycetous fungi. FEMS Microbiol Rev 2008;32:501–21. 10.1111/j.1574-6976.2008.00106.x.

[116] Tan T-C, Kracher D, Gandini R, Sygmund C, Kittl R, Haltrich D, et al. Structural basis for cellobiose dehydrogenase action during oxidative cellulose degradation. Nat Commun 2015;6:7542. 10.1038/ncomms8542.

[117] Zhang X-X, Gao J-S, Zhang L, Zhang C-W, Ma X-T, Zhang J. *Bacillus oryzisoli* sp. nov., isolated from rice rhizosphere. Int J Syst Evol Microbiol 2016;66:3432–6. 10.1099/ijsem.0.001215.

[118] Li S, Tang Y, Fang X, Qiao T, Han S, Zhu T. Whole-genome sequence of *Arthrinium phaeospermum*, a globally distributed pathogenic fungus. Genomics 2020;112:919–29. 10.1016/j.ygeno.2019.06.007.

[119] Khan SA, Hamayun M, Kim H, Yoon H, Seo J, Choo Y, et al. A new strain of *Arthrinium phaeospermum* isolated from *Carex kobomugi* Ohwi is capable of gibberellin production. Biotechnol Lett 2009;31:283–7. 10.1007/s10529-008-9862-7.

[120] Su L, Zhu H, Guo Y, Du X, Guo J, Zhang L, et al. *Lecanicillium coprophilum* (Cordycipitaceae, Hypocreales), a new species of fungus from the feces of *Marmota monax* in China. Phytotaxa 2019;387:55. 10.11646/phytotaxa.387.1.4.

[121] Nguyen TT, Le TN-G, Nguyen TH. First report of emerging fungal pathogens of *Cordyceps militaris* in Vietnam. Sci Rep 2023;13:17669. 10.1038/s41598-023-43951-9.

[122] Woolley VC, Teakle GR, Prince G, de Moor CH, Chandler D. Cordycepin, a metabolite of *Cordyceps militaris*, reduces immune-related gene expression in insects. J Invertebr Pathol 2020;177:107480. 10.1016/j.jip.2020.107480.

[123] Kato T, Nishimura K, Suparmin A, Ikeo K, Park EY. Effects of Cordycepin in *Cordyceps militaris* during Its Infection to Silkworm Larvae. Microorganisms 2021;9:681. 10.3390/microorganisms9040681.

[124] Vuong HQ, McFrederick QS. Comparative Genomics of Wild Bee and Flower Isolated *Lactobacillus* Reveals Potential Adaptation to the Bee Host. Genome Biol Evol 2019;11:2151–61. 10.1093/gbe/evz136.

[125] McFrederick QS, Vuong HQ, Rothman JA. *Lactobacillus micheneri* sp. nov., Lactobacillus timberlakei sp. nov. and Lactobacillus quenuiae sp. nov., lactic acid bacteria isolated from wild bees and flowers. Int J Syst Evol Microbiol 2018;68:1879–84. 10.1099/ijsem.0.002758.

[126] Liu Y, Qiao J-T, Yuan X-Z, Guo R-B, Qiu Y-L. *Hydrogenispora ethanolica* gen. nov., sp. nov., an anaerobic carbohydrate-fermenting bacterium from anaerobic sludge. Int J Syst Evol Microbiol 2014;64:1756–62. 10.1099/ijs.0.060186-0.

[127] Zhang X-X, Gao J-S, Zhang L, Zhang C-W, Ma X-T, Zhang J. *Bacillus oryzisoli* sp. nov., isolated from rice rhizosphere. Int J Syst Evol Microbiol 2016;66:3432–6. 10.1099/ijsem.0.001215.

[128] Ueki A, Ohtaki Y, Kaku N, Ueki K. Descriptions of *Anaerotaenia torta* gen. nov., sp. nov. and *Anaerocolumna cellulosilytica* gen. nov., sp. nov. isolated from a methanogenic reactor of cattle waste and reclassification of *Clostridium aminovalericum*, *Clostridium jejuense* and *Clostridium xylanovorans* as *Anaerocolumna* species. Int J Syst Evol Microbiol 2016;66:2936–43. 10.1099/ijsem.0.001123.

[129] Murawska A, Migdał P, Zajdel B, Popiela E, Roman A. The composition of red mason bee cocoons. J Apic Res 2022;61:227–32. 10.1080/00218839.2021.1988278.

[130] Hammer TJ, Moran NA. Links between metamorphosis and symbiosis in holometabolous insects. Philosophical Transactions of the Royal Society B: Biological Sciences 2019;374:20190068. 10.1098/rstb.2019.0068.

[131] Johnston PR, Rolff J. Host and Symbiont Jointly Control Gut Microbiota during Complete Metamorphosis. PLoS Pathog 2015;11:e1005246. 10.1371/journal.ppat.1005246.

[132] Poulsen M, Bot ANM, Currie CR, Nielsen MG, Boomsma JJ. Within-colony transmission and the cost of a mutualistic bacterium in the leaf-cutting ant *Acromyrmex octospinosus*. Funct Ecol 2003;17:260–9. 10.1046/j.1365-2435.2003.00726.x.

[133] Martinson VG, Moy J, Moran NA. Establishment of Characteristic Gut Bacteria during Development of the Honeybee Worker. Appl Environ Microbiol 2012;78:2830–40. 10.1128/AEM.07810-11.

[134] Koch H, Abrol DP, Li J, Schmid-Hempel P. Diversity and evolutionary patterns of bacterial gut associates of corbiculate bees. Mol Ecol 2013;22:2028–44. 10.1111/mec.12209.

[135] Powell JE, Martinson VG, Urban-Mead K, Moran NA. Routes of Acquisition of the Gut Microbiota of the Honey Bee *Apis mellifera*. Appl Environ Microbiol 2014;80:7378–87. 10.1128/AEM.01861-14.

[136] Kaltenpoth M, Goettler W, Koehler S, Strohm E. Life cycle and population dynamics of a protective insect symbiont reveal severe bottlenecks during vertical transmission. Evol Ecol 2010;24:463–77. 10.1007/s10682-009-9319-z.

[137] Kölsch G, Matz-Grund C, Pedersen B V. Ultrastructural and molecular characterization of endosymbionts of the reed beetle genus *Macroplea* (Chrysomelidae, Donaciinae), and proposal of “ Candidatus *Macropleicola appendiculatae*” and “ Candidatus Macropleicola muticae.” Can J Microbiol 2009;55:1250–60. 10.1139/W09-085.

[138] Buchner P. Endosymbiosis of animals with plant microorganisms. New York, NY: John Wiley & Sons 1965.

[139] Wolschin F, Hölldobler B, Gross R, Zientz E. Replication of the Endosymbiotic Bacterium *Blochmannia floridanus* Is Correlated with the Developmental and Reproductive Stages of Its Ant Host. Appl Environ Microbiol 2004;70:4096–102. 10.1128/AEM.70.7.4096-4102.2004.

[140] Tejerina MR, Cabana MJ, Enríquez PA, Benítez-Ahrendts MR, Fonseca MI. Bacterial Strains Isolated from Stingless Bee Workers Inhibit the Growth of *Apis mellifera* Pathogens. Curr Microbiol 2024;81:106. 10.1007/s00284-024-03618-8.

